# Anatomy of a pest control failure: Introgression of cytochrome P450 337B3 alleles from invasive old-world bollworm into native corn earworm (Lepidoptera: Noctuidae)

**DOI:** 10.1101/2024.03.22.584691

**Authors:** Marissa I. Nufer, Brad S. Coates, Craig A. Abel, Patrick O’Neill, Morgan McCracken, Devendra Jain, Calvin A. Pierce, James Glover, Tyler Towles, Gadi VP Reddy, Omaththage P. Perera

## Abstract

The establishment of invasive species populations can threaten the ecological balance in naïve habitats and impact agricultural production practices. *Helicoverpa armigera* (old-world bollworm, OWBW) and *H. zea* (corn earworm, CEW) were geographically separated prior to the 2013 report of OWBW invasion into South America. Introgression of OWBW-specific cytochrome P450 337B3 (CYP337B3) gene into CEW was repeatedly detected across South America and the Caribbean. Two hybrids were documented from Texas in 2019. In this study, screening insects collected in Olathe, Colorado, USA, where a failure of pyrethroids to control CEW damage to conventional sweetcorn in 2023 detected 28.6% of insects with the OWBW-specific CYP337B3 marker. Nucleotide sequencing of the CYP337B3 gene identified 73.1 and 26.9% of insects carried CYP337B3v2 and CYP337B3v6 alleles, respectively and 0.15 overall frequency of CYP337B3 alleles. Based on prior data for distinct phylogeographic origins of CYP337B3v2 and v6 alleles, our results indicate Olathe samples were derived from two different introductions; An uncertain source of the v6 allele that was initially reported in West Africa and possibly South American or Caribbean origin of the globally distributed v2 allele. One of the 1618 individuals screened also carried a ribosomal RNA internal transcribed spacer 1 (ITS1) derived from OWBW. Local selection pressures at the Olathe location imposed by repeated pyrethroid exposures are likely attributed to the prevalence of CYP337B3, where control practices hasten the accumulation of phenotypic resistance by adaptive introgression. Pyrethroid and other resistance factors carried by invasive OWBW may continue to impact CEW management tactics across the Americas.

## Introduction

Invasive insect species can disrupt naïve ecosystems, impact the health of native species, and introduce challenges to agricultural production (Venette and Hutchison 2021). An invasive species undergoes an initial establishment phase with varying levels of adaptation to new ecosystems and environmental conditions (Colautti et al. 2017). Ironically, invasive populations typically undergo a genetic bottleneck due to a small number of founders, where this reduced genetic diversity would reduce adaptive potential. Bottleneck effects can be alleviated through admixture following multiple introduction events from genetically distinct source populations, where increased genetic diversity in admixed populations can lead to higher fitness (Jaspers et al. 2021). Establishment can also be facilitated by interspecific hybridization with native species, which can transfer locally adapted alleles to invasive populations through a mechanism referred to as adaptive introgression (Arnold and Martin 2009, Jin et al. 2023). Conversely, native populations may also acquire alleles with a selective advantage from an invasive species (Burgarella et al. 2019)

The sister species *Helicoverpa armigera* (Hübner) (Lepidoptera: Noctuidae) and *H. zea* (Boddie) (Lepidoptera: Noctuidae) are morphologically similar but distinguishable via differences in male genitalia (Pouge 2004). The endemic range of *H. zea* spans the Americas, wherein larvae cause feeding damage to multiple high-value crops (Reay-Jones 2019). *Helicoverpa armigera* feeds on a broader range of host plants (Fitt 1989) and is historically distributed across Africa, Asia, Europe, and Oceania (Kitching and Rawlins 1998). Invasion and establishment of *H. armigera* into the Americas were first reported in Bahia, Goiás, and Mato Grosso states in Brazil in 2013 using morphological characters and the histology of male genitalia (Czepak et al. 2013). The presence of *H. armigera* was confirmed through analysis of mitochondrial DNA (Tay et al. 2013). Later, *H. armigera* was reported in other regions of Brazil and additional South American countries including Argentina, Colombia, Paraguay, Uruguay, and Venezuela (Specht et al. 2013, Arnemann et al. 2016, Arnemann et al. 2019, Riaz et al. 2021). Evidence suggests that these *H. armigera* populations resulted from multiple independent introductions (Tay et al. 2017, Arnemann et al. 2019, Tembrock et al. 2019). Laboratory experiments produced viable progeny from reciprocal crosses and backcrosses between *H. armigera* and *H. zea* (Laster and Sheng 1995), demonstrating incomplete barriers to reproduction. Genomic analyses of *Helicoverpa* populations detected evidence of hybridization between *H. armigera* and *H. zea* among field collections, and the introgression of *H. armigera* alleles into *H. zea* (Valencia-Montoya et al. 2020). These findings include the characterization of introgression of the *H. armigera* locus for a cytochrome P450, CYP337B3, associated with insecticide resistance into the *H. zea* genomic background (Walsh et al. 2018).

High mobility of migratory *Helicoverpa* and its modeling predict a high probability of *H. armigera* spreading and establishing in North America (Kriticos et al. 2015). *Helicoverpa armigera* was initially detected outside South America in Puerto Rico and Florida in 2014 and 2015, respectively (Smith 2014, Hayden and Brambila 2015, USDA-APHIS 2024). *Helicoverpa armigera* genome introgression was also predicted using genomic DNA sequence data from prior studies (Anderson et al. 2018) to characterize four *H. zea* samples collected from New York in 2017 as hybrids (Trujillo et al. 2024). The pest surveillance program of the USDA Animal and Plant Health Inspection Service (USDA APHIS) later detected several *H. armigera* adults near the Chicago O’Hare International Airport between 2019 and 2022, however, no individuals were detected in surrounding agricultural areas (USDA-APHIS 2024). Introgression between these two species was shown by the detection of the invasive *H. armigera-*specific CYP337B3 allele in two *H. zea* collected from a Texas population in 2019 (North et al. 2024). Our current study focuses on a collection of *Helicoverpa* moths from Olathe, Colorado, where a failure to control damage to conventional sweetcorn was reported following repeated pyrethroid insecticide applications in 2023. Noctuid pest management effort involved initial applications of fenvalerate and mid-season applications of methomyl, chlorantraniliprole, nucleopolyhedroviruses, and Spinetoram. Evaluations of increased carrier volumes for aerial applications, maximum labeled rate use per spray, maximum allowable frequency on repeat applications, and daily and every-other-day spraying regimes were all undertaken, using the aforementioned active ingredients. An additional active ingredient, methoxyfenozide, labeled for sweet corn and lepidopteran pests, was trialed under the direction of the product’s manufacturer to evaluate the potential for that material to be used via fixed-wing aircraft, with reduced carrier volume applications. However those trials proved ineffective at adding to the suppression of larval populations in the sweetcorn fields.

Currently a limited number of genetic markers are available to screen invasive *H. armigera.* Regardless, our analyses of species-specific genetic markers at ITS1 (Perera et al. 2015) and CYP337B3 loci (Joußen et al. 2012) detected *H. armigera* alleles in the Olathe samples. DNA from *H. zea* collected in 2006 from Mississippi and Texas was used as the reference. Our work constitutes the second report of *H. zea* carrying genes introgressed from *H. armigera* within the USA. Pyrethroid resistance conferred by the movement of the CYP337B3 allele into the *H. zea* population has implications for the continued efficacy of insecticidal control tactics used to manage this crop pest.

## Materials and Methods

Insects were collected using *Helicoverpa zea* pheromone lures in bucket traps (Trece, Adair, OK) at 53 locations near Olathe, CO, USA, during the summer of 2023 (**Supp. Table S1 and Supp. Fig. S1**). A sweetcorn grower near these traps reported that repeated applications of pyrethroid insecticide failed to control insect damage to the crop, and local scouting provided visual identification of surviving larval *H. zea*. Initial screening of insects was carried out by pooling a single leg from each adult male from each trap collection (or all the detached legs in a collection bag) and extracting DNA using a modified squish buffer containing 10 mM Tris-HCl, 0.25 mM EDTA, and 0.25 mM NaCl (Perera et al. 2015). Two 3-mm stainless steel beads were added to the tubes containing insect legs and homogenized using a bead beater (Biospec Products, Bartlesville, OK) for 1 min at the highest speed. Homogenates were centrifuged briefly and heated for 5 min in a heating block set to 100 °C, followed by centrifugation at 12,500 x g for 10 min. DNA was also extracted from *H. zea* collected in 2006 from ten counties in Texas (*n* = 200) and five counties in Mississippi (*n* = 100), and included in this study. For these, a small piece of thoracic tissue from five insects was pooled into one tube to extract DNA using 150 µl of squish buffer as described above.

Several strategies were used to identify invasive *H. armigera* in the samples and confirm the presence of introgressed genes. First, the legs of insects were pooled by collection date to perform ITS1 melt curve analysis that identifies *H. armigera* in bulk samples (Perera et al. 2015). Second, the *H. armigera* ITS1 gene was identified in a pooled set of samples, and the DNA of individual insects from that pool was PCR amplified and sequenced to confirm the presence of *H. armigera* ITS1 gene. Next, DNA from individual insects was PCR amplified with primers specific for the *H. armigera* CYP337B3 gene followed by verification of hemizygous or homozygous state of the CYP337B3 gene by PCR amplification of CYP337B1 and CYP337B2 genes using gene gene-specific primers. Lastly, longer PCR amplicons for CYP337B3 were generated and sequenced to obtain nucleotide sequences for the differentiation of alleles.

The supernatant from each extract was transferred to a new tube and used in the melt curve analysis (Perera et al. 2015). Briefly, 1 µl of each DNA pool was used in a 20 µl reaction containing final concentrations of 1X Fast SYBR Master mix (Life Technologies Corporation, Carlsbad, CA) 0.4 µM common forward primer 3373Ha_Hz_ITS1F for internal transcribed spacer 1 (ITS1) of ribosomal RNA gene, *H. armigera*-specific reverse primer 3374Ha_ITS1R, and 0.2 µM of *H. zea*-specific reverse primer 3377Hz_ITS1R (**Table 1**). DNA amplification and melt curve analysis were performed in an ABI7500 Fast real-time PCR instrument (Applied Biosystems, Foster City) with 95 °C initial denaturation for 30 seconds followed by 40 cycles of denaturation at 95 °C for 5 seconds and annealing and extension at 60 °C for 30 seconds. A melt curve step was performed after the amplification cycles to determine the presence of species-specific amplicons. Agarose gel electrophoresis was performed to visualize the amplicons as a verification step.

**Table 1.**
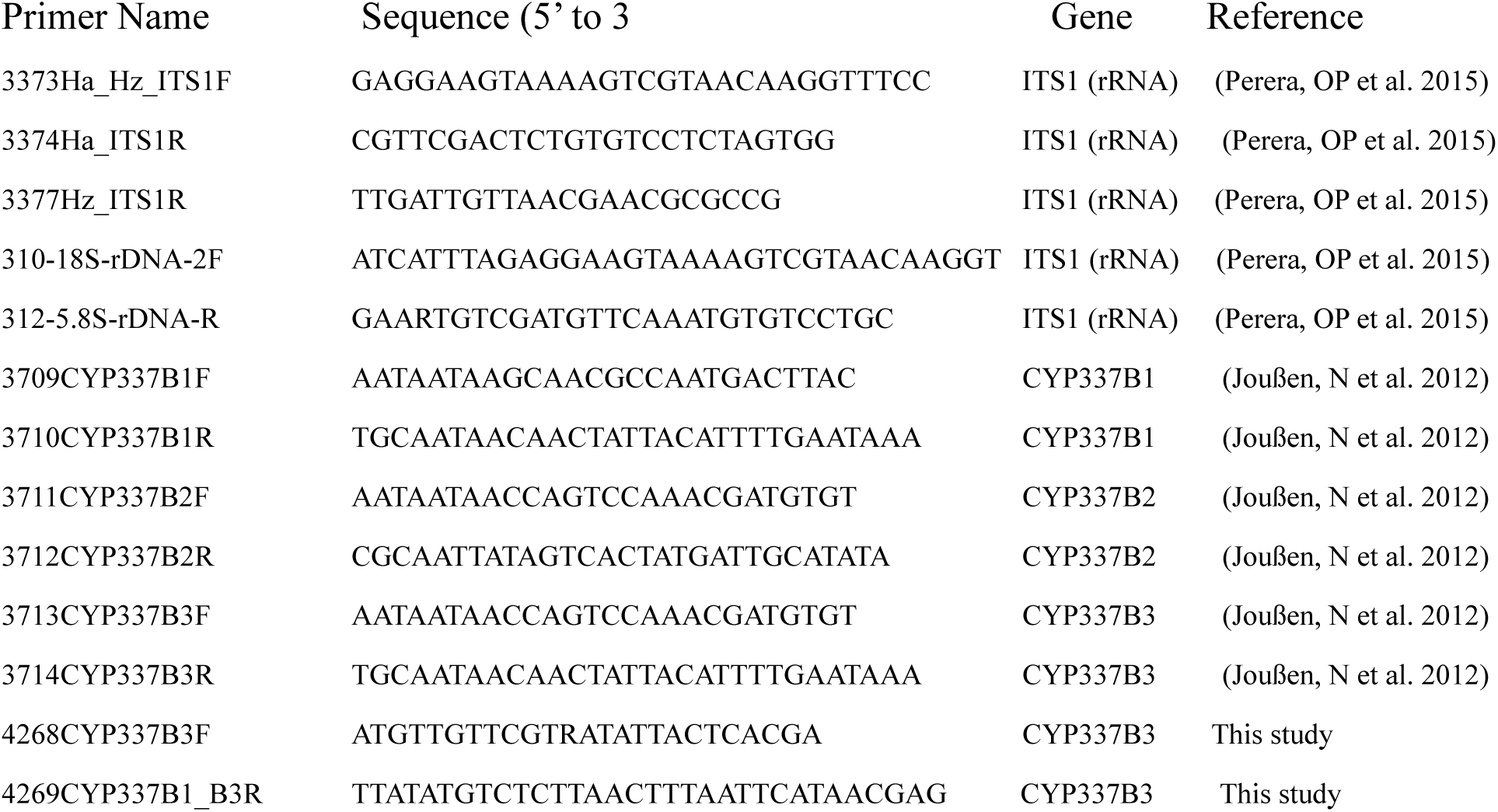
Primers used in PCR amplification and nucleotide sequencing of ribosomal RNA internal transcribed spacer 1 and cytochrome P450 337B, CYP337B3, from *Helicoverpa* samples collected from Olathe, Colorado, USA.

Full-length ITS1 was amplified from individual DNA using the primers 310-18S-rDNA-2F and 312-5.8S-rDNA-R (**Table 1**) in reactions containing 1X ThermoPol buffer (12.5 mM tricine, 42.5 mM KCl, 1.5 mM MgCl_2_, 6% dextran; New England BioLabs, Ipswich, MA), 0.32 mM dNTP mix, 0.4 µM of each primer, 1 unit of *Taq* polymerase, and 10–100 ng of genomic DNA in a 25 ml reaction. Amplification was performed in a PTC-200 thermal cycler (MJ Research, BioRad, Hercules, CA) with a 30 s initial denaturation step at 95 °C, 35 cycles of 15 s denaturation at 95 °C, 10 s annealing at 52 °C followed by a final extension of 5 min at 72 °C (Perera et al. 2015). Amplicons were purified using AmPure XP magnetic beads according to manufacturer instructions (Beckman Coulter, Brea, CA). Approximately 2 ng of amplicon from 82 separate pools was used as input for the Oxford Nanopore library construction reagent set SQK-LSK114 and the native DNA barcode kit SQK-NDB-96. The resulting 82 indexed libraries were sequenced in a FLO-MIN114.001 flow cell following the manufacturer’s instructions (Oxford Nanopore Technologies, New York, NY). Purified ITS1 DNA from sample 61e was also cloned into TA cloning vector (Invitrogen, Carlsbad, CA) and transformed into DH5α competent cells. Plasmid DNA prepared from 24 colonies were screened for the presence of *H. armigera-* and *H. zea-*specific amplicons as described previously. Nucleotide sequences of clones positive for ITS1 specific for each species were obtained by Sanger sequencing. DNA Sequences were analyzed using CLC Genome WorkBench v22.02 (Qiagen, Redwood City, CA).

Phylogenetic analysis was performed using Molecular Evolutionary Genetic Analysis (MEGA) 11 (Tamura et al. 2021) based on a multiple sequence alignment of the ITS1 region from *H. armigera*, *H. zea,* and *H. assulta* (Guenée) (KT342378.1, KT343380.1, and KT343379.1, respectively), and our Sanger read data for Olathe, CO, individual 61e. Relationships were estimated using the Maximum Likelihood (ML) method (Nei and Kumar 2000, Beerli and Felsenstein 2001) applying the Kimura-2-Parameter model of sequence evolution (Kimura 1980), that implemented a discrete Gamma shape parameter and node support inferred from 10,000 bootstrap pseudo replications (Felsenstein 1985) of the aligned data. The tree was rooted by the *Chloridea* (*Heliothis*) *virescens* (Fabricius) ITS1 (GenBank accession KT342279.1)

A short fragment of the cytochrome P450 337B3 (CYP337B3) gene was amplified from Olathe, CO, Texas and Mississippi samples using gene-specific primers 3713CYP337B3F and 3714CYP337B3R (**Table 1**) following the protocol described in Joußen et al. (2012). Briefly, the 25µl amplification reactions contained 1x standard Taq polymerase buffer (10 mM Tris-HCl, 50 mM KCl, pH 8.3) (New England Biolabs, Ipswich, MA), 2.5 mM MgCl_2_, 0.4 µM dNTP mix, 0.4 µM of each primer, and 0.025 units of Taq polymerase. A thermal cycling profile with a 15 s initial denaturation step at 95 °C, 35 cycles of 10 s denaturation at 95 °C, 10 s annealing at 53°C, an extension of 30 sec at 72 °C followed by a final extension of 5 min at 72 °C.

Amplicons were resolved by 1.0% agarose gel electrophoresis to identify insects positive for the CYP337B3 gene. For each insect identified as positive for CYP337B3, the full-length CYP337B3 gene was then amplified with primers 4268CYP337B3F and 4269CYP337B1_B3R. PCR amplification reaction and thermal cycling parameters were the same as those used for amplification of the short fragment of CYP337B3, except for an annealing temperature of 50 °C and extension time increased to 2.75 min. Amplicons were purified using AmPure XP beads (Beckman Coulter) prior to use in sequencing reactions. Barcoded Oxford Nanopore libraries were constructed and sequenced as described above for full-length CYP337B3 gene sequences from 143 insects. CYP337B3 allele versions v1 through v8 (JQ995292.1, KJ636466.1, KM675665.1, KM675666.1, KX958388.1, KX958389.1, KX958390.1, and KX958391.1, respectively), CYP337B1v1, and CYP337B2v2 (JQ995291) were used as references for subsequent alignment of Oxford Nanopore amplicon sequences. Evolutionary relationships of the CYP337B3 alleles from Olathe, CO population and previously published CYP337B genes was inferred in an ML tree constructed using MEGA 11 as described above.

The CYP337B1 and CYP337B2 genes were subsequently PCR amplified from all CYP337B3 positive DNA samples in reactions with the primers for CYP337B1 and CYP337B2 genes using the primer pairs 3709CYP337B1F/3710CYP337B1R and 3711CYP337B2F/3712CYP337B2R, respectively (Table 1). PCR conditions for amplification of CYP337B1 and CYP337B2 were identical to those used to amplify the CYP337B3 gene, and resulting amplicons were resolved by electrophoresis in 1.0% agarose gels.

## Results

Initial screening of 1618 insects collected from Olathe, CO in 2023 occurred within 82 pools by PCR with *H. armigera*-specific ITS1 primers, where an amplicon was generated from a single pool, pool 61. This indicated at least one *H. armigera* or a hybrid insect in pool 61 carried an *H. armigera* ITS1 region sequence. In contrast, none were detected in the Texas or Mississippi samples collected in 2006. Analysis of Oxford Nanopore sequence data from pool 61 also identified ITS1 sequences closely resembling *H. armigera* ITS1. Screening of DNA from individual insects from pool 61 identified one insect (61e) from which both *H. armigera-* and *H. zea*-specific ITS1 amplicons were produced. Subsequent alignment found that the ribosomal RNA (rRNA) gene sequence of insect 61e contained both *H. zea* rRNA repeats and a variant of *H. armigera* rRNA repeats. Specifically, this was shown from Oxford Nanopore sequencing and Sanger sequencing of cloned full-length ITS1 PCR products from 61e (**Fig. 1**) where insertion/deletion (indel) mutations were shown in the *H. armigera* compared to *H. zea* ITS1 references sequences at positions 138-167, 401-403, and 431-433. Sequence data from sample 61e showed two variants, with indels in each identical to respective *H. armigera-* and H. zea ITS1 reference sequences. However, there were 23 single nucleotide polymorphisms in the *H. armigera-*like ITS1 sequence from sample 61e that were identical to the *H. zea* ITS1 reference. Phylogenetic reconstruction of ITS1 region showed clustering of the two ITS region variants from Olathe, CO individual 61e with corresponding *H. armigera* (KT342378.1) and *H. zea* references (KT343380.1; **Fig. 2**). ITS1 nucleotide sequences from the Olathe, CO sample 61e were deposited in the GenBank database under accession numbers PP456864.1 (Olathe_61e_Ha_ITS1), and PP456865.1 (Olathe_61e_Hz_ITS1).

**Figure 1.**
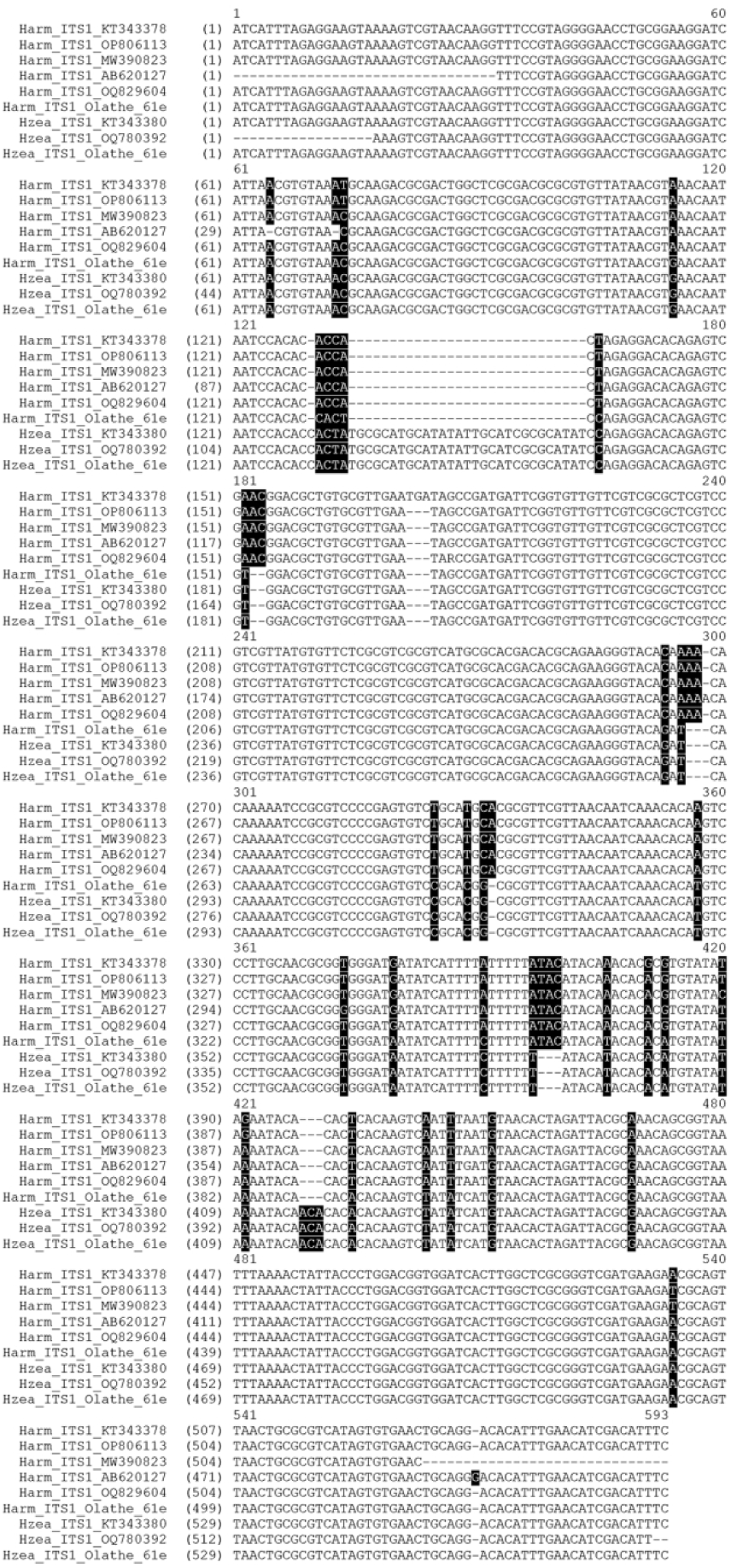
Alignment of reference ITS1 sequences of *Helicoverpa armigera* (Ha) and *H. zea* (Hz) with ITS1 sequences obtained from Olathe, CO sample 61e. Nucleotides identical in both *H. armigera* and *H. zea* ITS1 sequences are shown in black. Olathe_61e_Ha_ITS1 sequence was highly similar to *H. zea* ITS1 sequence except for the *H. armigera*-specific ITS1 sequences. Differences between ITS1 sequences from the two *Helicoverpa* species are shown by white text with black background. Alignment gaps are indicated by a hyphen (-).

**Figure 2.**
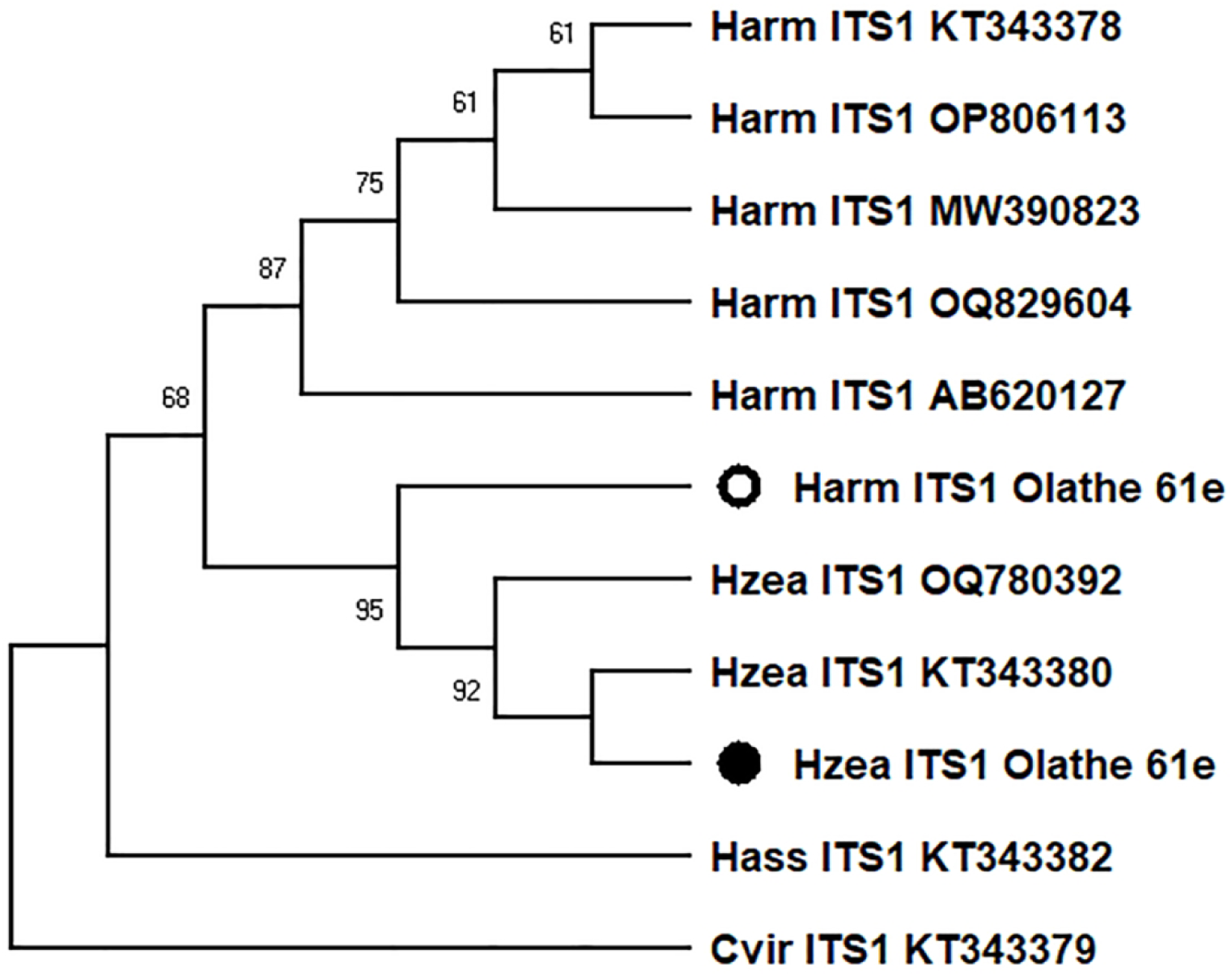
The evolutionary history was inferred among internal transcribed spacer 1 (ITS1) regions using Maximum Likelihood based on the Kimura 2-parameter model of sequence evolution (Kimura, 1980). The phylogeny among ITS1 regions among Helicovera armigera (Harm), *H. zea* (Hzea) and *H. assulta* (Hass) was rooted by the *Chloridea (Heliothis) virescens* (Cvir) ITS1 (GenBank accession provided at branch tips), based on a 601 nucleotide-long consensus alignment of 11 nucleotide sequences. The bootstrap consensus at each node was inferred from 10,000 replicates (Felsenstein, 1985) and partitions with less than 50% bootstrap support are collapsed. Neighbor-Join and BioNJ algorithms were applied to a matrix of pairwise distances estimated using the Maximum Composite Likelihood (MCL) approach and the topology with superior log likelihood value was selected to obtain the initial tree(s). A discrete Gamma distribution was used to model evolutionary rate differences among sites (5 categories (+G, parameter = 0.3073)). Evolutionary analyses were conducted in MEGA11 (Tamura et al. 2021).

Ribosomal RNA units in eukaryotes are arranged as head-to-tail tandem repeats at one or more loci (Nei and Rooney 2005, Naidoo et al. 2013). One study indicated that the rRNA gene copy number in animals is highly variable, ranging from 39 in the mosquito *Sabethes cyaneus* (Fabricius) to 19,300 in the salamander *Amphiuma means* Garden in Smith (Prokopowich et al. 2003). Although precise copy number estimates are not available for noctuid moths, the silk moth *Bombyx mori* (L.) has approximately 240 copies of rRNA genes (Prokopowich et al. 2003). In the reference *H. zea* genome assembly, ilHelZeax1.1 (RefSeq assembly GCF_022581195.2; (Stahlke et al. 2023), BioProject: PRJNA818699), one complete set of rRNA repeats and one 18S rRNA subunit are annotated at one end of the chromosome 30 (Accession: NC_061481.1; XR_006986104.1, XR_006986099.1, and XR_006986099.1). Similarly, one rRNA repeat unit is annotated at one end of chromosome 30 in *H. armigera* (Accession: OZ005796.1; BioProject: PRJEB72184). Collapse of identical rRNA repeat units during the assembly of the genome may have reduced the number of repeats in the final assembly.

Tandem CYP337B genes (paralogs) are located in chromosome 2 of the *H.* zea reference genome (Stahlke et al. 2023) and the orthologous chromosome 15 of the *H. armigera* genome (Zhang et al. 2022). Different alleles of CYP337B3 resulted from independent recombination events that joined exon 1 of the CYP337B2 gene with exon 2 of the CYP337B1 gene (Joußen et al. 2012, Rasool et al. 2014, Walsh et al. 2018) (**Supp. Fig. S2a**). Therefore, wild-type tandem genes CYP337B1 and CYP337B2 and recombinant CYP337B3 genes in *H. zea* can occur in five different genotypic configurations: homozygous *H. zea* CYP337B1/CYP337B2 genes (**Supp. Fig. S2b**), heterozygous *H. armigera* and *H. zea* CYP337B1/CYP337B2 genes (**Supp. Fig. S2c**), hemizygous *H. armigera* CYP337B1/CYP337B2 and CYP337B3 genes (**Supp. Fig. S2d**), hemizygous *H. zea* CYP337B1/CYP337B2 and CYP337B3 genes (**Supp. Fig. S2e**), and homozygous CYP337B3 genes (**Supp. Fig. S2f**). Insects homozygous or hemizygous for introgressed CYP337B3 gene can be identified using gene-specific primers (Joußen et al. 2012). PCR screening of insects from the Olathe, CO, collection from 2023 with primers 3713CYP337B3F and 3714CYP337B3R identified 243 of 850 (28.6%) as PCR positive and carrying at least one copy of a CYP337B3 gene. Screening all 243 samples positive for the CYP337B3 gene with primers for CYP337B1 and CYP337B2 genes indicated that 12 samples were negative for both wild-type genes and homozygous for the CYP337B3 gene. Remaining insects were hemizygous with wild-type CYP337B1/CYP337B2 genes on one chromosome and CYP337B3 gene on the other chromosome. Therefore, the overall allele frequency of CYP337B3 in Olathe, CO, population of *H. zea* estimated from 12 homozygous and 231 hemizygous insects was 0.15 (255/1700). The estimated frequencies of CYP337B3 homozygotes, hemizygotes, and wild-type homozygotes in the Olathe, CO, population were 0.0225, 0.2550, and 0.7225, respectively. The observed frequency of CYP337B3 homozygotes (0.0141) was less than the expectation under the Hardy-Weinberg equilibrium whereas the observed frequency of hemizygotes was higher.

None of the samples in our study from Texas or Mississippi collected in 2006 generated amplicons for the *H. armigera* CYP337B3 gene. Oxford Nanopore sequencing and mapping of individual reads from 113 full-length amplicons and 28 short amplicons from 141 CYP337B3 positive insects from Olathe, CO, predicted 100 and 99.3% nucleotide identity to the reference genomic sequences of CYP337B3 alleles v2 (KJ636466.1) and v6 (KX 958389.1), respectively. The subsequent consensus built from mapped reads (coverage > 200-fold) indicated that CYP337B3 alleles v2 (accession#: PP461124) and v6 (accession#: PP461125) were present among individuals from Olathe, CO. Approximately 26.9% of the analyzed samples (38 out of 141) had CYP337B3v6 while the remaining were from the CYP337B3v2 allele.

Four of the samples identified with the CYP337B3v2 allele also contained 5.3 to 7.6% Oxford Nanopore sequence reads that mapped to the CYP337B3v6 allele. The percentages of CYP337B3v6 reads identified in these four samples were significantly lower than the 50% reads expected from a heterozygous genotype with v2 and v6 alleles. Whether these reads resulted from cross-contamination of samples during collection in traps and shipping in bags as bulk samples, errors in base calling of barcodes during Nanopore sequencing, or PCR bias is unknown.

Alignment of the mRNA and genomic sequences of CYP337B1v1, CYP337B2v2, and CYP337B3 alleles v1-v8, including two consensus CYP337B3v2 and CYP337B3v6 alleles sequences found in the Olathe, CO, showed Olathe CYP337B3v2 genomic and mRNA nucleotide sequences were nearly 100% identical to the reference CYP337B3v2 (KJ636466.1) sequence (**Supp. Figs. S3 and S4**). Specifically, there was one synonymous T to C substitution in the coding regions between the reference CYP337B3v2 allele (KJ636466.1) and the CYP337B3v2 gene variant present in the Olathe, CO population (**Supp. Fig. S5**). Alignments also showed that CYP337B3v6 genomic DNA and mRNA sequences were 99.4% and 98.9% identical to the reference CYP337B3v6 (KX 958389.1) allele, respectively. There were 16 SNPs between the reference CYP337B3v6 allele from Uganda (KX958389.1) and the v6 allele variant found in Olathe, CO (**Supp. Fig. S6**), but only a single mutation, an A to T transversion at the nucleotide position 232, resulted in a conservative isoleucine to leucine substitution at the amino acid position 78 (I78L). The consensus ML tree indicated that predicted CYP337B3v2 and v6 alleles from the Olathe, CO population clustered with corresponding reference alleles with node support from ≥ 99% of the bootstrap iterations (**Fig. 3**).

**Figure 3.**
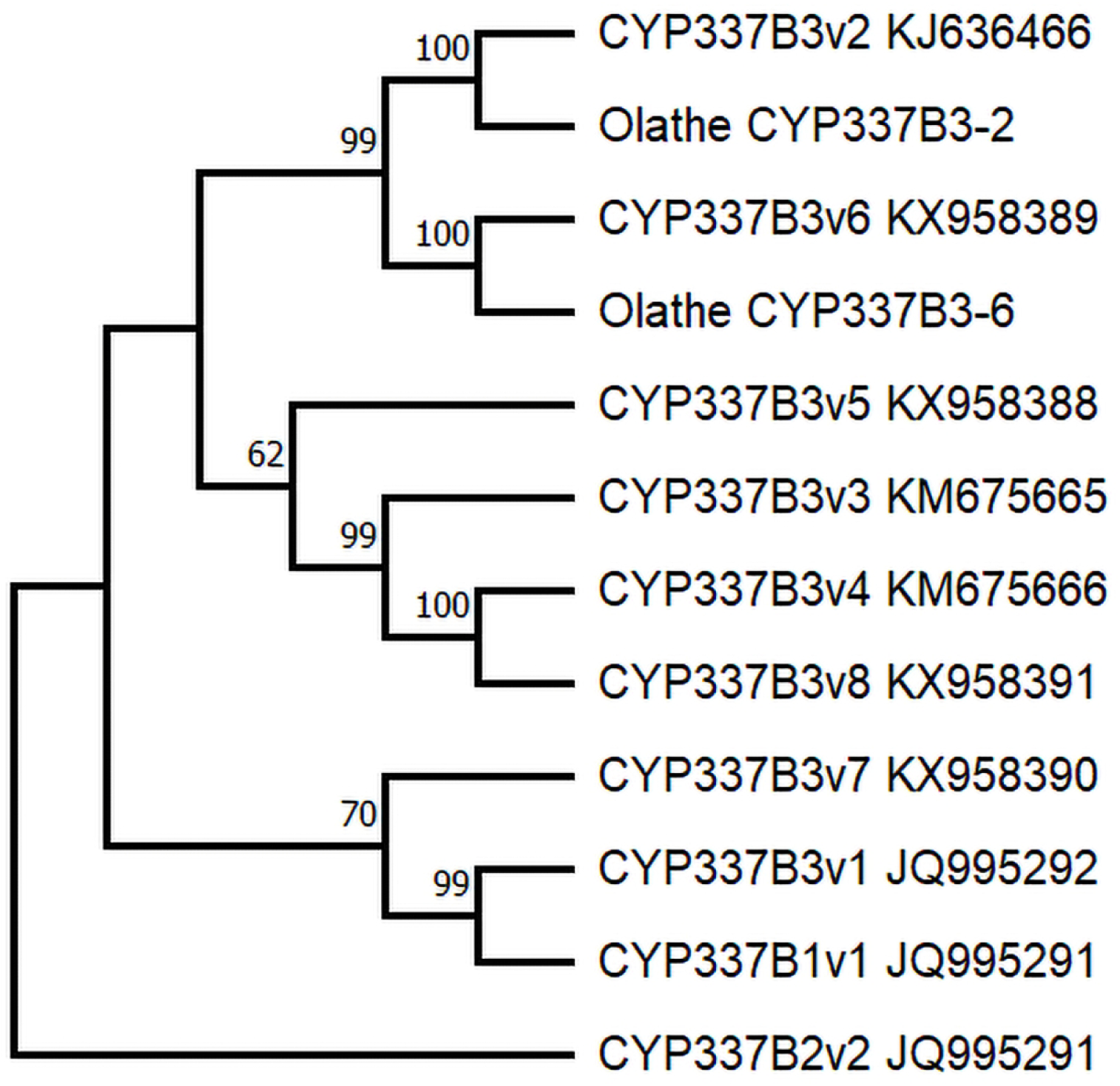
The evolutionary history was inferred among cytochrome P450 337B orthologs, CYP3B1-3, from *Helicoverpa armigera* and *H. zea* by using Maximum Likelihood and based on the Kimura 2-parameter model of sequence evolution (Kimura, 1980) with 10,000 bootstrap replicates (Felsenstein, 1985) as implemented in Mega 11 (Tamura et al. 2021). Relationships of alleles from the Olathe, CO population are shown with respect to reference sequences for CYP337B1, CYP337B2 and CYP337B3 paralogs (GenBank accessions include at respective branch tips) and based on 3605 positions in a consensus sequence alignment. Neighbor-Join and BioNJ algorithms were applied to a matrix of pairwise distances estimated using the Maximum Composite Likelihood (MCL) approach and the topology with superior log likelihood value was selected to obtain the initial tree(s). A discrete Gamma distribution was used to model evolutionary rate differences among sites (5 categories (+G, parameter = 0.3794) and the tree with the highest log likelihood (−9917.03) is shown.

## Discussion

The chimeric CYP337B3 gene associated with the functional cause of pyrethroid resistance in *H. armigera* evolved from an unequal crossover between paralogs CYP337B1 and CYP337B2 and had not evolved natively in *H. armigera* (Joußen et al. 2012, Rasool et al. 2014). The *H. armigera* CYP337B3 gene confers resistance to class II pyrethroid insecticides, including cypermethrin, deltamethrin, and fenvalerate (Rasool et al. 2014, Durigan et al. 2017). Evidence of hybridization between invasive *H. armigera* and native *H. zea* was previously reported in South America (Tay et al. 2013, Anderson et al. 2018), wherein the introgression of CYP337B3 into sympatric populations was detected (Valencia-Montoya et al. 2020, Ivey and Hillier 2023). The initial report of hybridization in North America detected CYP337B3 in two *H. zea* collected in Texas in 2019 that had no described or documented history of pyrethroid exposures (North et al. 2024). In this study, we found that approximately 28.6% of the adult *H. zea* collected in 2023 near Olathe, CO, where repeated pyrethroid applications by growers failed to control larval feeding damage to sweetcorn during the same growing season, carried one of two different alleles, CYP337B3v2 or CYP337B3v6, at the CYP337B3 locus. This CYP337B locus of *H. armigera* exists in homozygous, heterozygous, and hemizygous states for the CYP337B1/CYP337B2 pair and recombinant CYP337B3 (Walsh et al. 2018). Invasive *H. armigera* populations in the Americas may have contributed recombinant CYP337B3 to *H. zea* (Valencia-Montoya et al. 2020, North et al. 2024), which is detectable using a gene-and species-specific marker (Joußen et al. 2012). However, *H. armigera*-derived CYP337B1/CYP337B2 cannot be detected in hybrid *H. zea* using the PCR screening methods in this or prior studies. Evaluation remains limited to detecting alleles of hybrid *Helicoverpa* via the CYP337B3 gene and thus possibly underestimates the proportion of introgression. Out of 243 insects identified to be positive for the CYP337B3 gene, only 12 were homozygous, and the remaining insects were hemizygous at the CYP337B locus with wild-type CYP337B1/CYP337B2 gene pair from *H. zea* or *H. armigera*.

Reference *H. zea* populations from Texas and Mississippi collected in 2006 prior to the documented *H. armigera* invasion of South America in 2013 or North America in 2019 (North et al. 2024, USDA-APHIS 2024) did not contain the CYP337B3 gene. Our results thus suggest that the *H. armigera* invasion into the United States occurred after 2006 or was previously undetectable due to low frequency during its initial establishment. Regardless, a more comprehensive spatial and temporal screening of archived samples would be needed to approximate the time(s) and the location(s) of the introduction. Previous population genetic studies postulated that the CYP337B3 locus may be under strong selection in Brazil due to the association between the high frequency of this gene and failures of pyrethroid insecticides to control damage among field populations (Durigan et al. 2017, Valencia-Montoya et al. 2020). Pyrethroid resistance due to knock-down resistance (*kdr*) mutations in the voltage-gated sodium channel (*vgsc*) and significantly increased expression of cytochrome P450 6B (*cyp6b*) genes was previously reported in the *H. zea* populations in the USA (Pietrantonio et al. 2007, Hopkins and Pietrantonio 2010, Hopkins et al. 2011). In contrast, *kdr* mutations were not reported among pyrethroid-resistant *H. armigera* populations in China, whereas the CYP337B3 gene was fixed among all samples tested (Ni et al. 2023).

The above lines of evidence, along with introgression of genome regions flanking CYP337B (Durigan et al. 2017, Valencia-Montoya et al. 2020, North et al. 2023), support the premise that the chimeric CYP337B3 detected in the Americas originated from invasive *H. armigera*. Olathe is in western Colorado, where a discontinuous cropping landscape, punctuated by expanses of non-arable land and high mountain elevations, exists. Thus, the detection of the *H. armigera* CYP337B3 in *H. zea* from such remote and isolated fields may be considered not surprising even given the known mobility (Westbrook and López 2010, Jones et al. 2019) and high gene flow of *H. zea* within the continent (Seymour et al. 2016, Perera et al. 2020). Our findings, along with that of two hybrid *H. zea* with the CYP337B3 gene found in samples collected in 2019 from Texas (North et al. 2023) and the dispersal behavior of this species might suggest hybrids exist in the regions of the southwestern United States. CYP337B3-positive insects identified in samples collected in New York, Texas, and Virginia from 2016 to 2023 (Perera, unpublished) most likely indicate the presence of hybrid *H. zea* with genes introgressed from *H. armigera* within the continent. Instance of continuous and repeated use of pyrethroid insecticides in the United States could provide high selection pressures, as previously hypothesized in South America, to facilitate adaptive introgression of the CYP337B3 gene (Durigan et al. 2017, Valencia-Montoya et al. 2020). Such a scenario might lead to transient increases in CYP337B3 frequency among local populations that could become homogenized due to high gene flow in the *H. zea* population. This admixture of locally adapted phenotypes may effectively remediate impacts on agricultural production practices through an overall reduction in resistance allele frequency in a landscape if persistent and widespread selective pressure is lacking or contributes to the spread and accumulation of resistance across the population. The latter possible outcome would depend on the prevalence of pyrethroid use by growers, be modulated by the implementation of integrated pest management (IPM) practices, and be impacted by the interaction of ecological, genetic, and other factors. Furthermore, pyrethroid resistance developed previously among *H. zea* in the United States (Abd-Elghafar et al. 1993, Jacobson et al. 2009) prior to this recent introgression of CYP337B3. Thus, evaluating phenotypic levels of pyrethroid resistance conferred by *H. zea kdr* and CYP6B mechanisms compared to those by *H. armigera*-derived CYP337B3 alleles may provide an estimate of any selective advantages attained by introgression. Regardless, in contrast to adaptive introgression resulting in the transfer of locally adapted alleles from indigenous populations of genetically-compatible species to invasive species (Edelman and Mallet 2021, Jin et al. 2023), our results and prior evidence suggest the reverse scenario exists in *Helicoverpa* due to the influence of selection by pyrethroid insecticide use.

It was postulated that all allelic variants of CYP337B3 formed independently in different geographic regions (Walsh et al. 2018). In Olathe, CO, population, the CYP337B3v2 allele was present in 73.1% of the hybrid insects, while the remaining hybrid insects had the CYP337B3 v6 allele. The high-throughput sequencing of CYP337B3 amplicons interestingly did not indicate the presence of CYP337B3v2/CYP337B3v6 heterozygotes among all 141 samples sequenced. Assuming the increase in metabolic capacity of homozygotes, the high prevalence of these alleles may be anticipated at locations such as Olathe, CO, where selection pressure was imposed following repeated exposures to pyrethroid applications. Regardless, mating between lineages carrying different haplotypes at the CYP337B3 locus would predict the formation of heterozygotes in the population unless admixture is suppressed due to recency of contact or yet undescribed barriers to gene flow exist. Our data indicate that the frequency of observed homozygotes is less than the estimated homozygote frequency. Additional research will be necessary to understand the fitness of different configurations of CYP337B genotypes.

The CYP337B3v2 allele was first reported in Pakistan and associated with an approximately 7-fold increase in resistance to cypermethrin (Rasool et al. 2014), and subsequently in Australia, China, and the African and European continents (Han et al. 2015, Walsh et al. 2018). In contrast, the CYP337v6 allele has only thus far been reported in Uganda (Walsh et al. 2018), and further description of any pyrethroid resistance phenotype it confers is currently lacking. In a previous study of *Helicoverpa*, almost all *H. armigera* analyzed from Brazil (*n* = 92) were reported homozygous for the CYP337B3v2 allele. At the same time, one insect was heterozygous for CYP337B3v2/ CYP337B3v5 alleles and found one of 59 *H. zea* carried the CYP337B3v2 allele (Walsh et al. 2018). Thus, the source population(s) for the introgressed CYP337B3v2 allele in Olathe, CO may be the South American *H. armigera* or hybrid *H. zea-armigera* populations that migrated or were transported northward, but an independent introduction source of CYP337B3 into North America cannot be ruled out.

Our study is the first to report the African-derived CYP337B3v6 allele in the Americas, providing evidence that a proportion of individuals in the Olathe population may be derived directly or indirectly from an African source population. It is possible that previous studies on South American populations of *H. zea* did not detect this allele due to very low frequencies on that continent. However, it should be noted that 26.9% of the CYP337B3 alleles observed in this study were CYP337B3v6 alleles, but this higher frequency could have resulted from genetic drift following a genetic bottleneck or selection upon introduction to North America. Therefore, it is not possible to determine if this allele was introduced directly from an African or South American population until additional genomic data are obtained. Although the point source(s) of the CYP337B3 alleles detected in Olathe, CO remains uncertain, a comparison of CYP337B3 flanking genomic regions between Olathe, CO and prior introgressed hybrids detected in North America (North et al. 2024) or *H. armigera* detected near O’Hare airport (USDA-APHIS 2024) might provide for such prediction. The sixteen nucleotide substitutions present between the CYP337B3v6 allele from Olathe, CO, and the v6 allele from Uganda may also point to an origin other than the African continent for CYP337B3v6 allele found in Olathe, CO. Although conserved amino acid substitutions such as I78L found in CYP337B3v6 allele from Olathe, CO, generally do not affect the function of a protein, alteration of function have been observed in some isoleucine to leucine substitutions (Sitbon et al. 1991, Brown et al. 2002, Christoffers et al. 2002). Therefore, the implications of this I78L substitution on resistance to pyrethroids remain unknown and in need of future empirical assessments. Nevertheless, the origins of two CYP337B3 alleles detected among *H. zea* from Olathe, CO, in this study indicate that at least two different hybridization events occurred between *H. armigera* and *H. zea* in the Americas, and the CYP337B3v6 allele could represent the signature of one or more new *H. armigera* introductions into the Americas.

The *H. armigera* rRNA ITS1 sequence identified in one insect (61e) in the Olathe, CO population further confirms introgression from *H. armigera* into *H. zea* through hybridization. We inferred that orthologous rRNA genes are located only in chromosome 30, because rRNA genes are not found on other chromosomes in the genome assemblies of either *Helicoverpa* species. It is conceivable that the crossover of orthologous chromosome regions in a hybrid progeny of *H. zea* and *H. armigera* could result in the introgression of some of the rRNA repeats into the *H. zea* genome. Regardless, several repetitive gene families in prokaryotes and eukaryotes are known to undergo concerted evolution, which functions to limit variation between repeat units (Coen et al. 1982, Mullins and Fultz 1989, Benedict et al. 1996, Liao 2000, Bettencourt and Feder 2002, Rooney and Ward 2005, Peel et al. 2006, Ganley and Kobayashi 2007, Naidoo et al. 2013). This insect 61e may have had rRNA gene repeats introgressed from *H. armigera* during a crossover event between *H. zea* and *H. armigera* haplotypes and in the process of gene conversion and repeated backcrossing to *H. zea.* However, it is not possible to determine the structure of the rRNA repeats in this insect and the origin of the sequences resembling ITS1 of both *H. armigera* and *H. zea* without haplotype phased sequence data from the segment of chromosome 30 containing the rRNA repeats. Concerted evolution would predict that the single nucleotide polymorphisms (SNP) specific to *H. armigera* ITS1 or the *H. zea* ITS1 would be converted over time, and unconverted variants may thus be indicative of how recent that introgression of the rRNA genes occurred. It should be noted that sample 61e was not positive for the CYP337B3 gene, indicating it may have originated from a hybridization event independent of those that led to the introgression of CYP337B1v2 and v6 alleles or the loci segregated independently in the population as expected for genes that have no known level of linkage disequilibrium. Thus, due to the lack of linkage between the ITS region and CYP337B loci, recombination in a backcross likely disentangled these two *H. armigera*-derived loci.

Invasive species are a threat to agricultural production in the United States and worldwide (Coates et al. 2015). A scenario of adaptive introgression facilitated by the transfer of a novel gene from an invasive to a native insect, as predicted for the *Helicoverpa* pest complex, provides a unique opportunity to study evolution and selection in real-time. Specifically, our evidence supports the admixture of previously geographically distinct introgressed CYP337B3v2 and CYP337B3V6 alleles into a single population. However, any hybrid vigor or advantageous phenotypic effects conferred by a unique combination of two pyrethroid resistance alleles in the North American population remains to be determined. The prevalence of these alleles across the continent remains unknown, high gene flow between North American *H. zea* populations (Perera et al. 2020) predicts these genes could spread rapidly across the continent. The CYP337B3 gene is currently used as a genetic marker to indicate the extent of gene transfer between these two species but likely underestimates actual introgression rates due to inability to detect *H. armigera*-derived CYP337B1/CYP337B2 haplotypes. The transfer of other genes with selective advantage, such as those conferring resistance to *Bacillus thuringiensis* insecticidal proteins, remains undetermined. The overall impacts of genes derived from invasive *H. armigera* on grower production practices remain uncertain, but continued selection for pest adaptions to practices that rely on insecticidal agents may be predicted to drive introgression of associated resistance alleles into *H. zea*. This may necessitate the development of alternative management strategies and the adoption of IPM practices.

## Acknowledgment

Authors thank Fanny Liu, USDA-ARS Biological Control Pests Research Unit, Stoneville, MS and Tina Ward and Miriam Lopez, USDA-ARS Corn Insects& Crop Genetics Research Unit, Ames, IA, for assistance with Sanger nucleotide sequencing. Mention of trade names or commercial products in this article is solely for the purpose of providing specific information and does not imply recommendation or endorsement by the US Department of Agriculture. USDA is an equal opportunity provider and employer.

## Author contributions

Conceptualization: MIN, OPP, and BSC. Methodology: MIN, OPP, CAP, DJ, and JG. Investigation: MIN, OPP, CAP, BSC, CAA, and DJ; Formal Analysis: MIN, OPP, BSC, CAA, and JG; Writing - Original Draft: MIN and OPP. Writing - Review & Editing: All authors. Resources: PON, MC, TT, and GVPR; Supervision: OPP.

---

**Table S1.** Trap numbers, GPS coordinates and the number of *Helicoverpa zea* samples used for screening CYP337B and ITS1 genes in this study.

| Trap Number | GPS Coordinates | Number of *H. zea* analyzed | Trap Number | GPS Coordinates | Number of *H. zea* analyzed |
| --- | --- | --- | --- | --- | --- |
| 132 | -108.011098, 38.562165 | 1 | 169 | -107.971621, 38.498372 | 13 |
| 134 | -108.015825, 38.547827 | 4 | 211 | -108.091077, 38.650081 | 12 |
| 135 | -107.916943, 38.489266 | 6 | 212 | -108.080870, 38.664141 | 14 |
| 136 | -108.011098, 38.562165 | 1 | 213 | -108.077449, 38.683565 | 17 |
| 137 | -108.054434, 38.584557 | 6 | 214 | -108.078346, 38.704736 | 12 |
| 138 | -108.015825, 38.547827 | 8 | 215 | -108.067737, 38.639066 | 29 |
| 141 | -108.068547, 38.649694 | 16 | 216 | -108.072320, 38.704661 | 24 |
| 142 | -108.059322, 38.60264 | 5 | 217 | -108.091077, 38.650081 | 25 |
| 145 | -107.909639, 38.487477 | 6 | 218 | -108.077434, 38.632005 | 74 |
| 146 | -108.073293, 38.591758 | 11 | 219 | -108.074783, 38.671722 | 62 |
| 149 | -108.016373, 38.540598 | 8 | 220 | -108.071173, 38.631818 | 46 |
| 150 | -108.016991, 38.551457 | 24 | 221 | -108.091077, 38.650081 | 60 |
| 151 | -108.053704, 38.610456 | 11 | 222 | -108.101899, 38.709269 | 105 |
| 152 | -108.111687, 38.657228 | 13 | 311 | -108.001276, 38.544657 | 2 |
| 153 | -108.044431, 38.646102 | 73 | 312 | -108.099386, 38.635561 | 28 |
| 154 | -108.016991, 38.551457 | 14 | 313 | -108.002041, 38.533634 | 7 |
| 155 | -107.960479, 38.489219 | 21 | 314 | -108.083571, 38.697376 | 12 |
| 156 | -108.050124, 38.584472 | 13 | 315 | -108.085612, 38.695765 | 15 |
| 157 | -108.023570, 38.558725 | 25 | 317 | -107.999232, 38.533403 | 9 |
| 158 | -108.023570, 38.558725 | 20 | 318 | -108.101954, 38.701580 | 22 |
| 159 | -108.040675, 38.575491 | 35 | 319 | -108.097406, 38.649842 | 2 |
| 160 | -107.990482, 38.539044 | 23 | 320 | -108.097406, 38.649842 | 4 |
| 161 | -108.040675, 38.575491 | 34 | 321 | -108.099386, 38.635561 | 14 |
| 162 | -107.950764, 38.474691 | 25 | 322 | -108.076984, 38.660800 | 30 |
| 163 | -107.914918, 38.487321 | 66 | 323 | -108.113262, 38.719266 | 66 |
| 164 | -107.960479, 38.489219 | 16 | 324 | -108.077128, 38.638438 | 81 |
| 165 | -107.970694, 38.496597 | 75 | 325 | -107.951346, 38.518697 | 37 |
| 166 | -107.954929, 38.551128 | 8 | 326 | -108.002959, 38.547599 | 92 |
| 167 | -107.954929, 38.551128 | 15 | CSU 1 | -108.070247, 38.660805 | 64 |
| 168 | -107.971621, 38.498372 | 20 | CSU 2 | -107.973397, 38.492867 | 37 |

**Figure S1.**
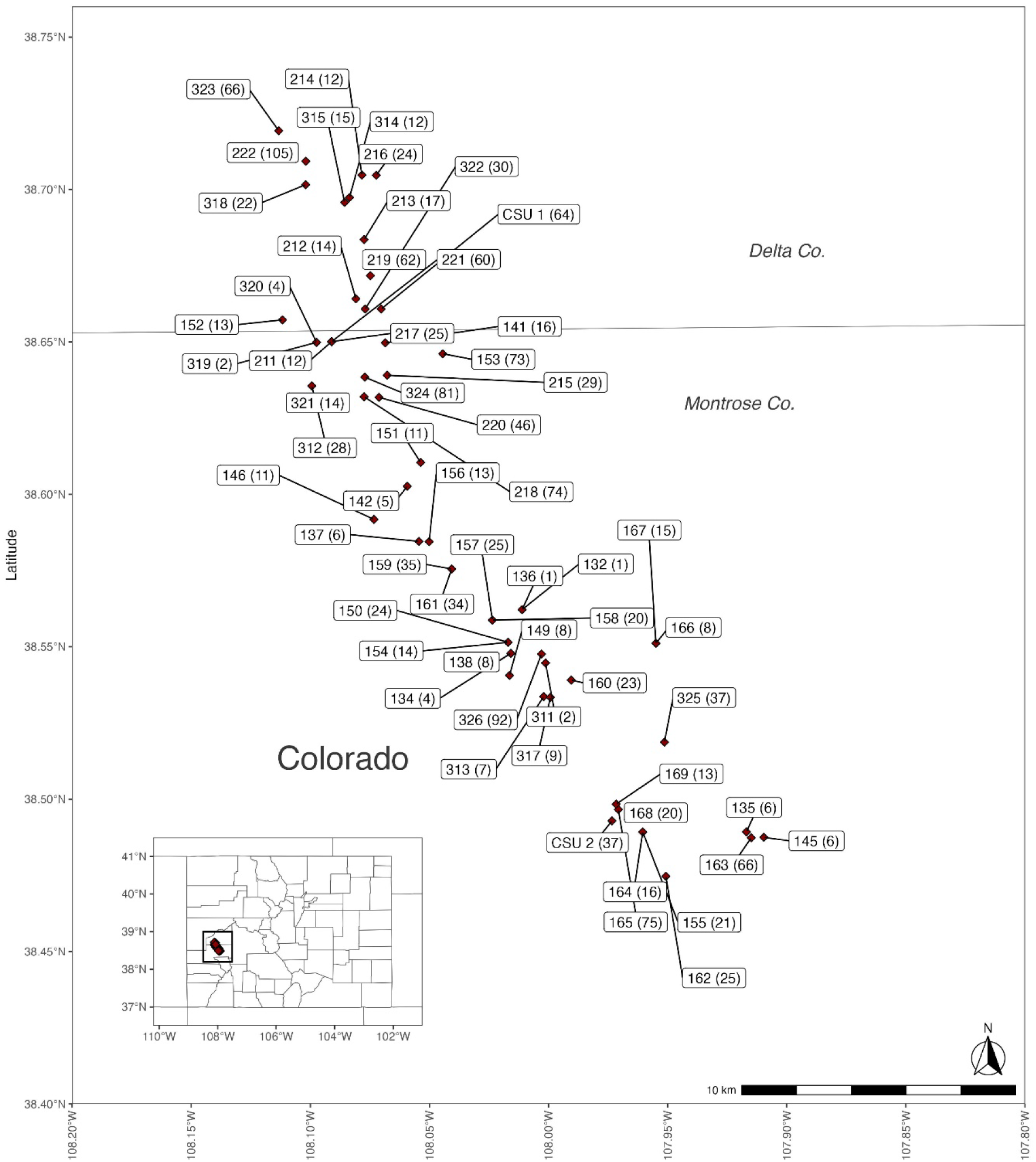
Locations of pheromone-lured traps used for collection of *Helicoverpa zea* in Olathe, Colorado, USA. Number of insects analyzed from each trap is indicated in parenthesis next to the trap number. R version 4.3.1 and the packages cowplot v. 1.1.3 (Wilke 2024), ggrepel v. 0.9.5 (Slowikowski 2024), ggsflabel v. 0.0.1 (Yutani 2024), ggspatial v. 1.1.9 (Dunnington 2023), maps v. 3.4.2 (Richard A. Becker, Ray Brownrigg. Enhancements by Thomas P Minka, and Deckmyn. 2023), rnaturalearth v. 1.0.1 (Massicotte and South 2023), rnaturalearthdata v. 1.0.0 (South, Michael, and Massicotte 2024), sf v. 1.0.16 (Pebesma 2018; Pebesma and Bivand 2023), tidyverse v. 2.0.0 (Wickham et al. 2019) were used in generating the site map using the data provided in the Supp. Table S1.

**Figure S2.**
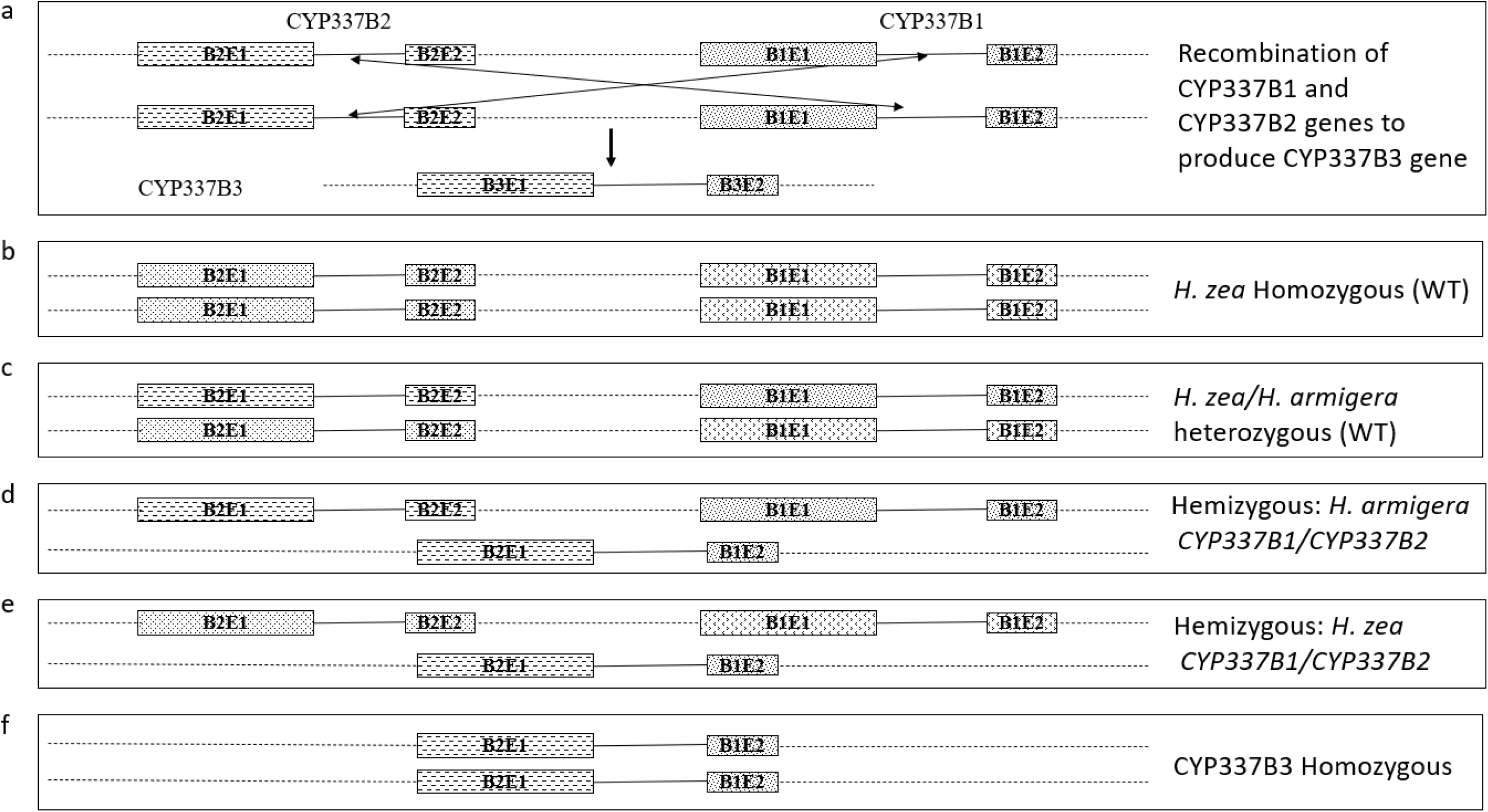
Schematic representation of CYP337B genes and potential genotypic configurations of CYP337B1/CYP337B2 genes and CYP337B3 genes in a population of *Helicoverpa zea* with CYP337B genes introgressed from *H. armigera.* a). unequal crossover between CYP337B1 and CYP337B2 genes to produce CYP337B3 gene. b). homozygous CYP337B1/CYP337B2 genes originated from wild type *H. zea.* c). heterozygotes of CYP337B1/CYP337B2 with each chromosome originating from *H. armigera* and *H. zea*. d). CYP337B3 hemizygotes with *H. armigera* CYP337B1/CYP337B2 genes. e). CYP337B3 hemizygotes with *H. zea* CYP337B1/CYP337B2 genes. f). CYP337B3 homozygotes. Allelic variants of different genes are not shown. B2E1: exon 1 of CYP337B2; B2E2: exon 2 of CYP337B2; B1E1: exon 1 of CYP337B1; B1E2: exon 2 of CYP337B1.

**Figure S3.**
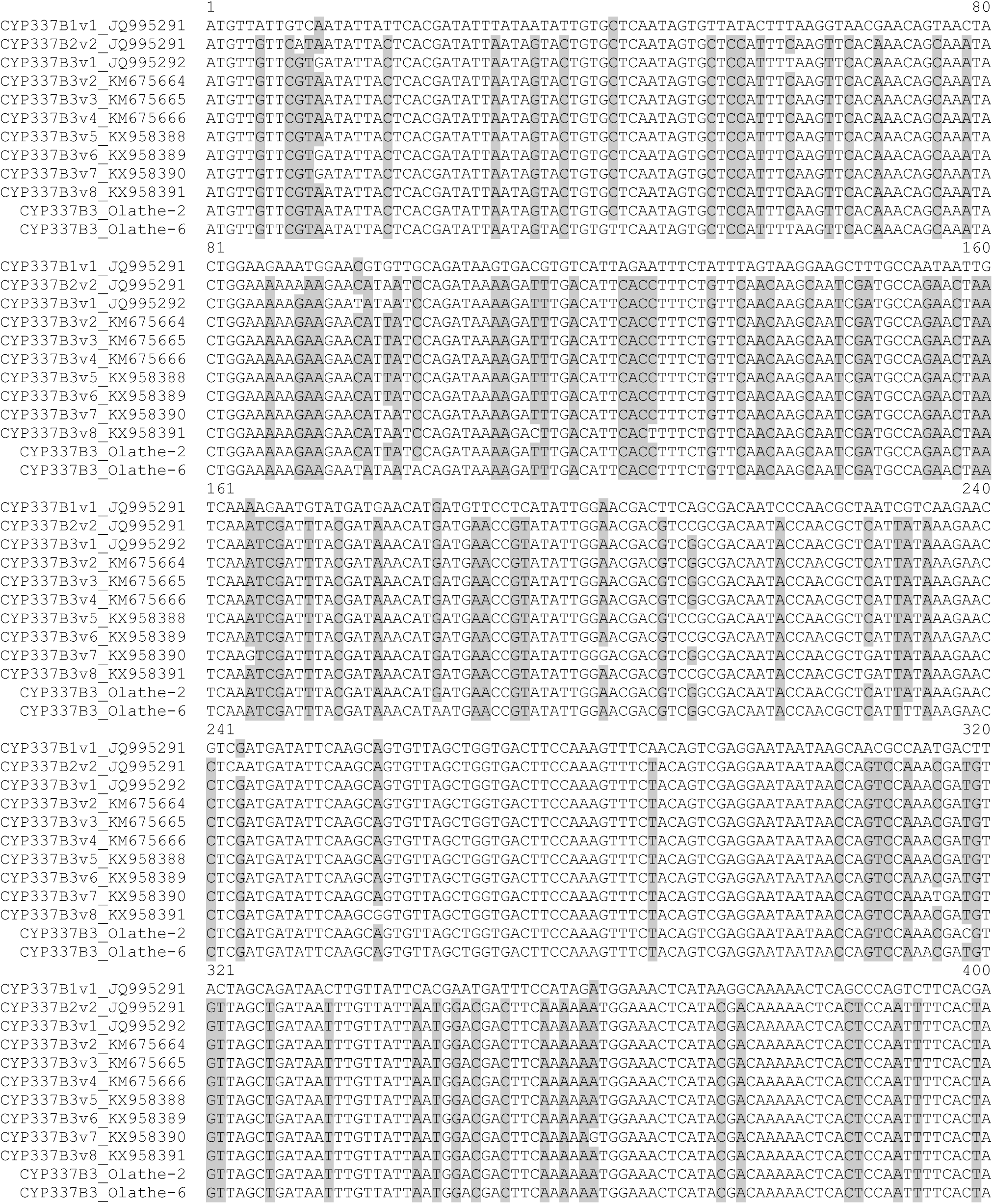
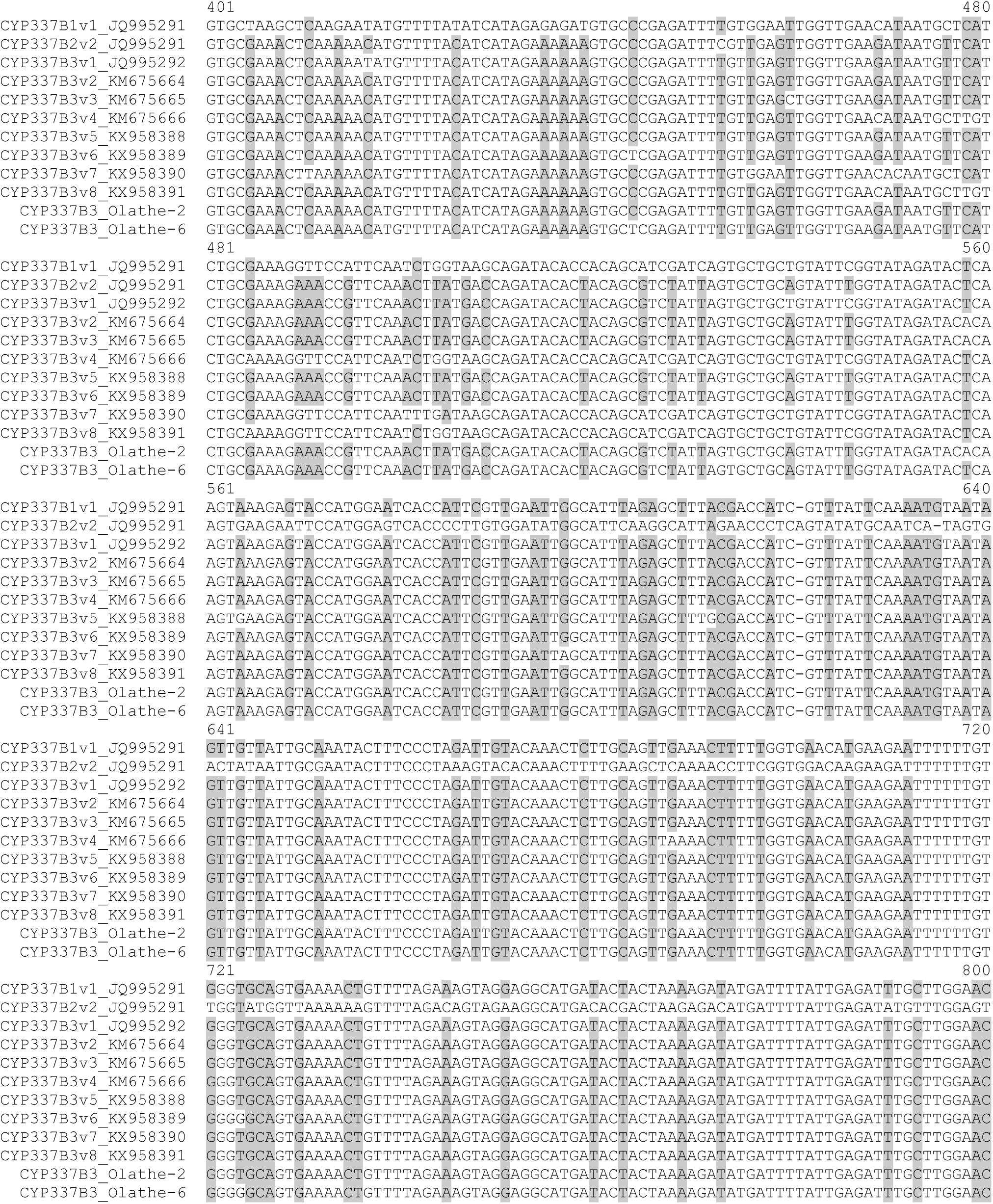
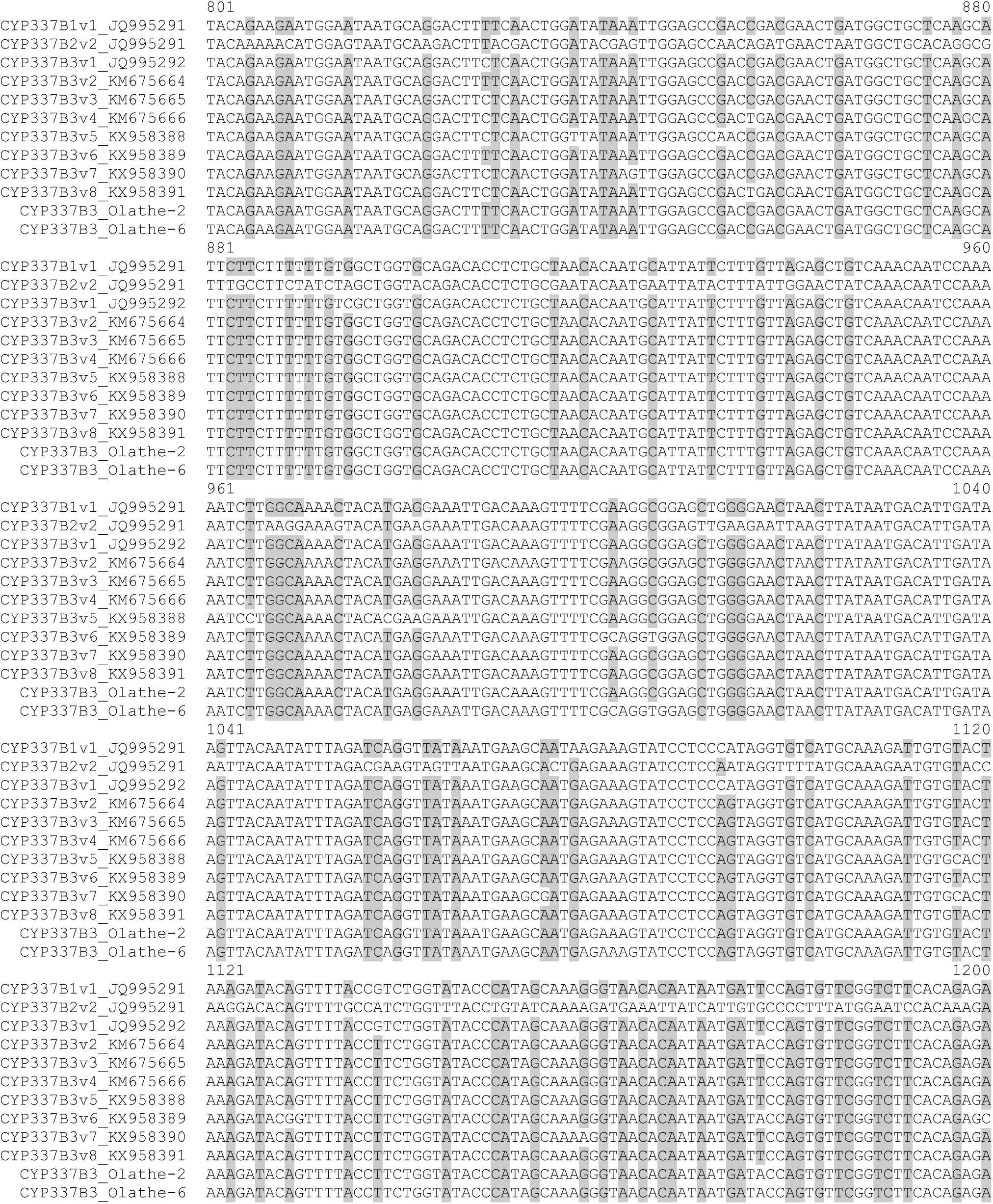
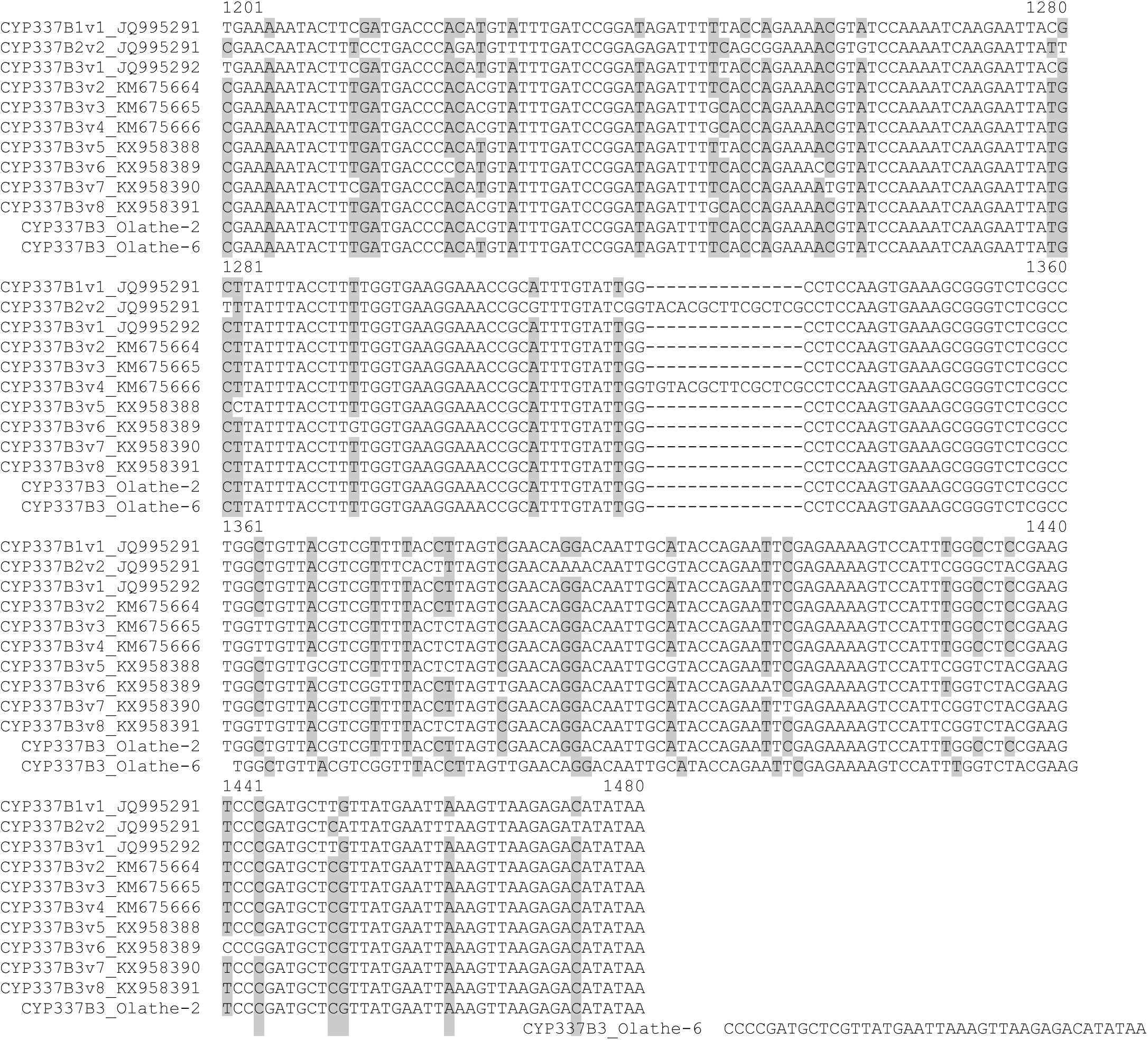
Alignment of the mRNA sequences of cytochrome P450 337B (CYP337B) genes CYP337B1v1, CYP337B2v2, and CYP337B3 alleles v1 through v8 with mRNA sequences extracted from CYP337B3 alleles identified from Olathe, CO population of *Helicoverpa zea*. Identical and conserved nucleotide positions are shown in black text and nucleotide polymorphisms, insertions, and deletions are shown with grey background. Alignment gaps are indicated with a hyphen (-).

**Figure S4.**
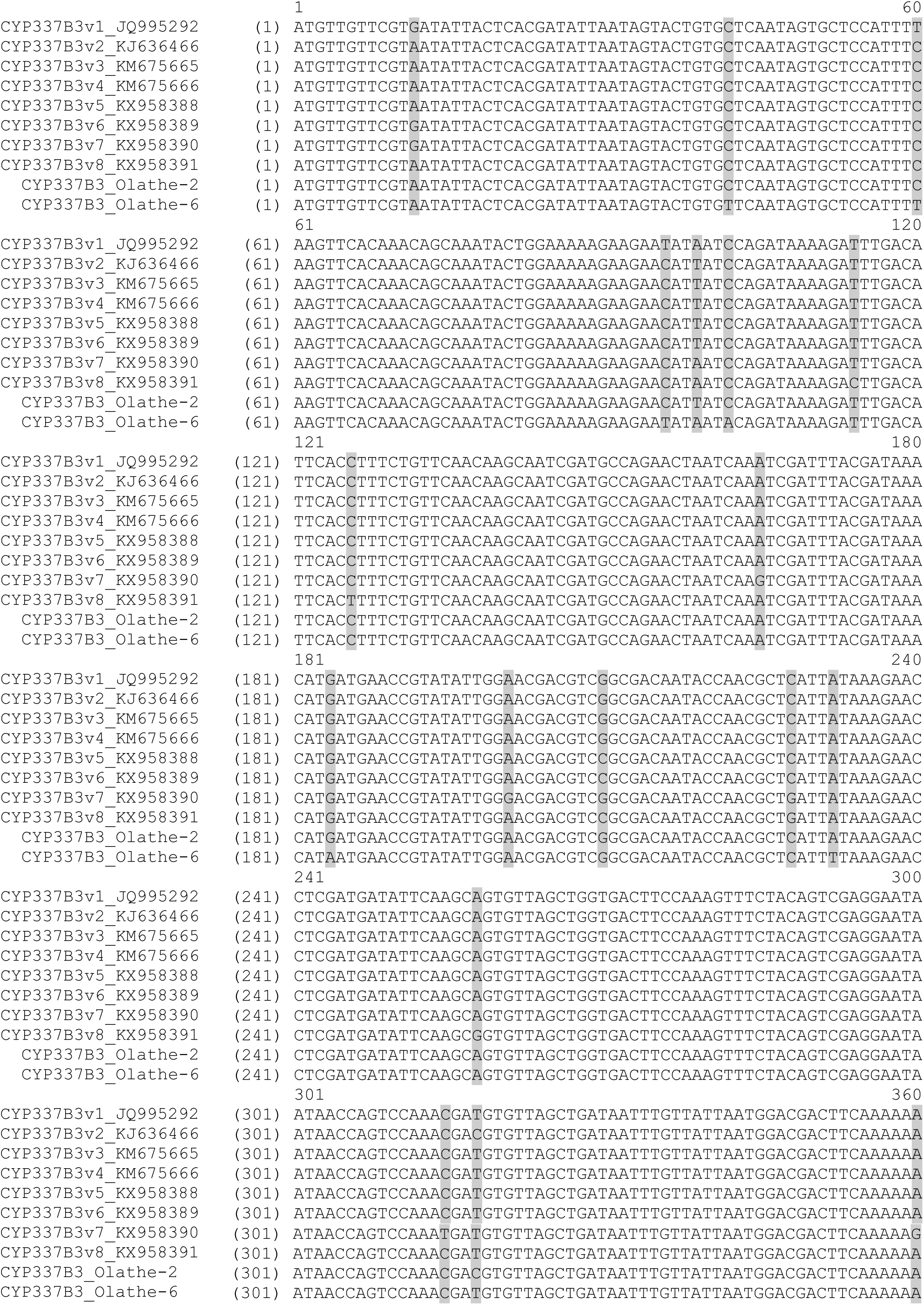
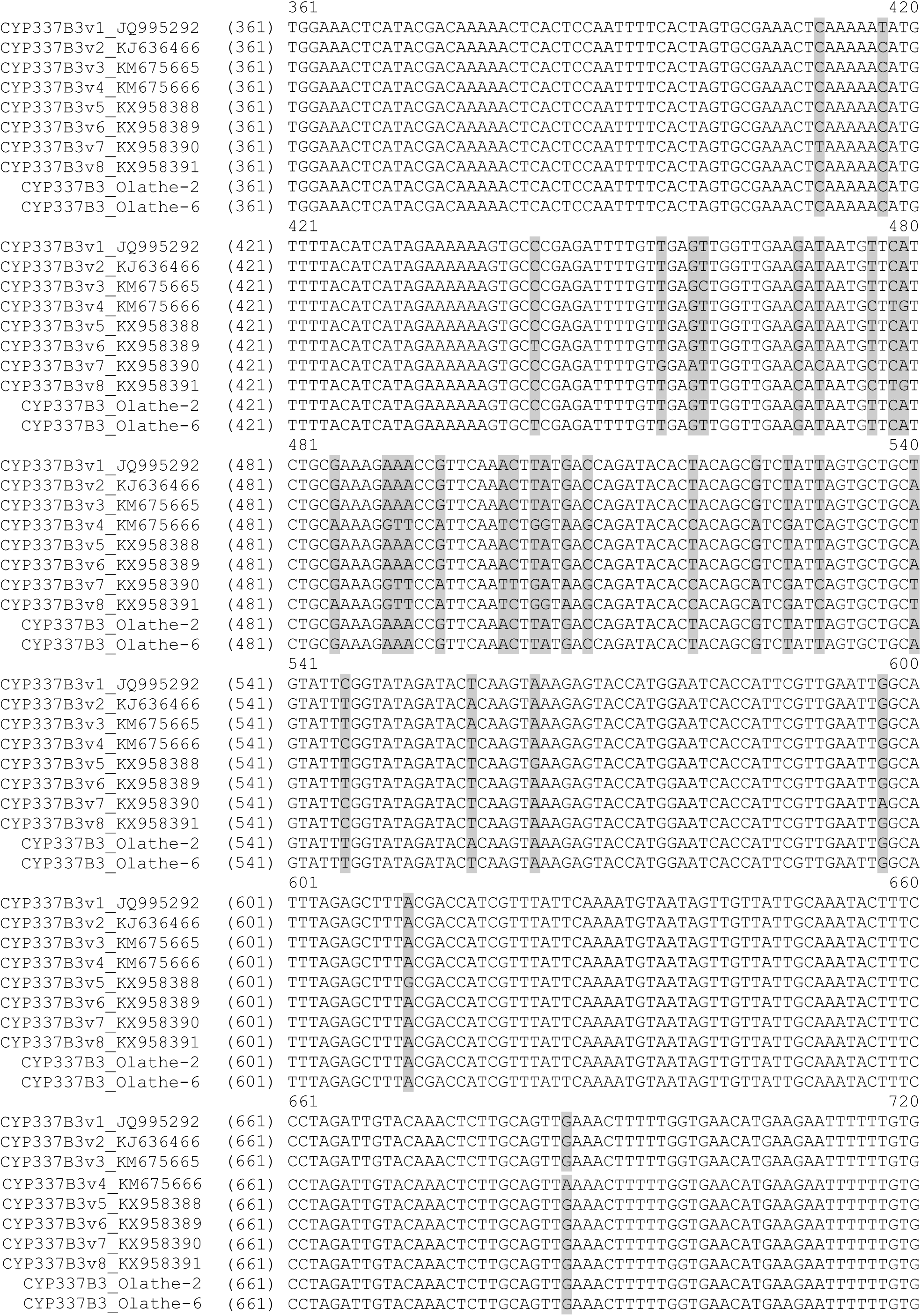
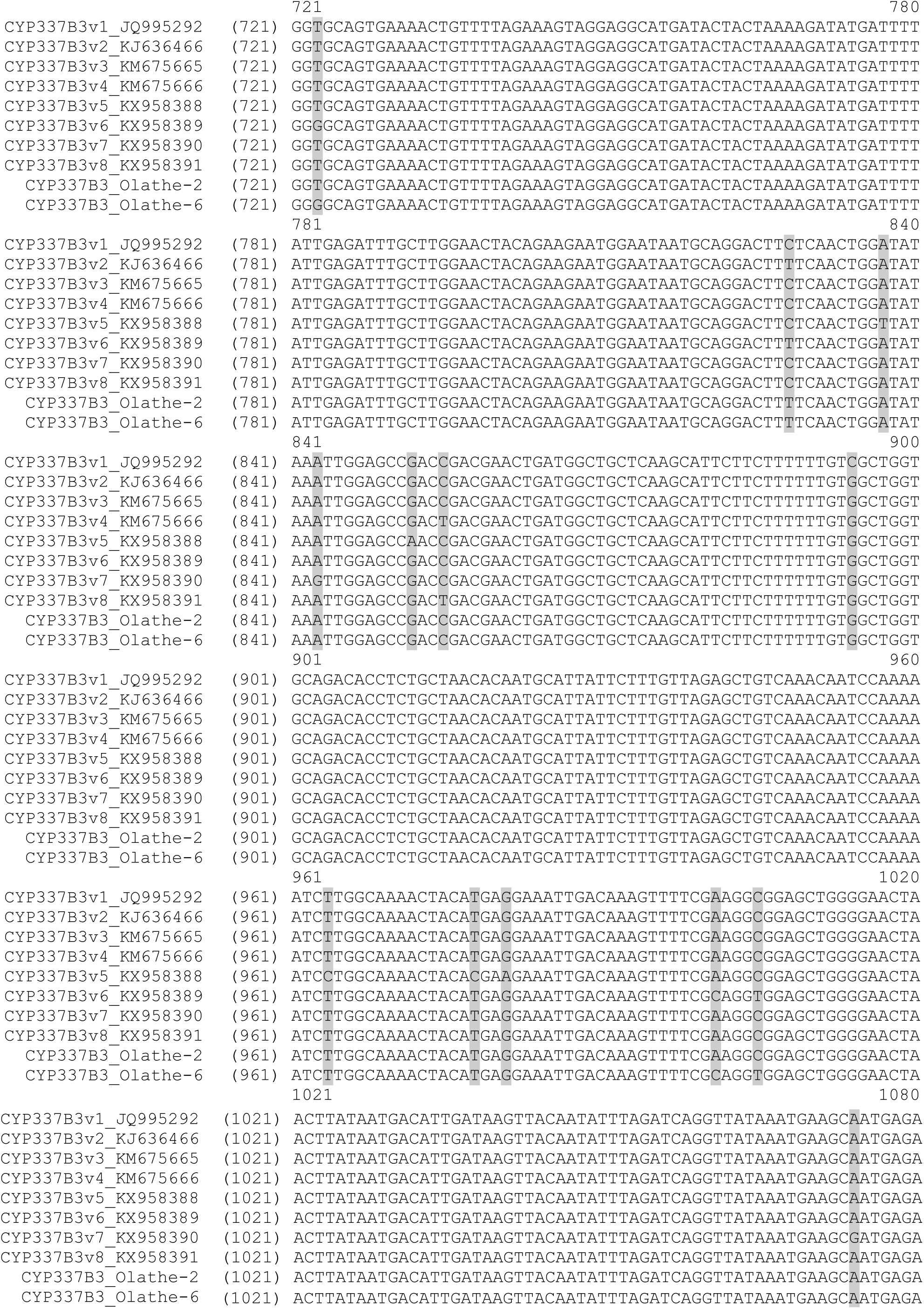
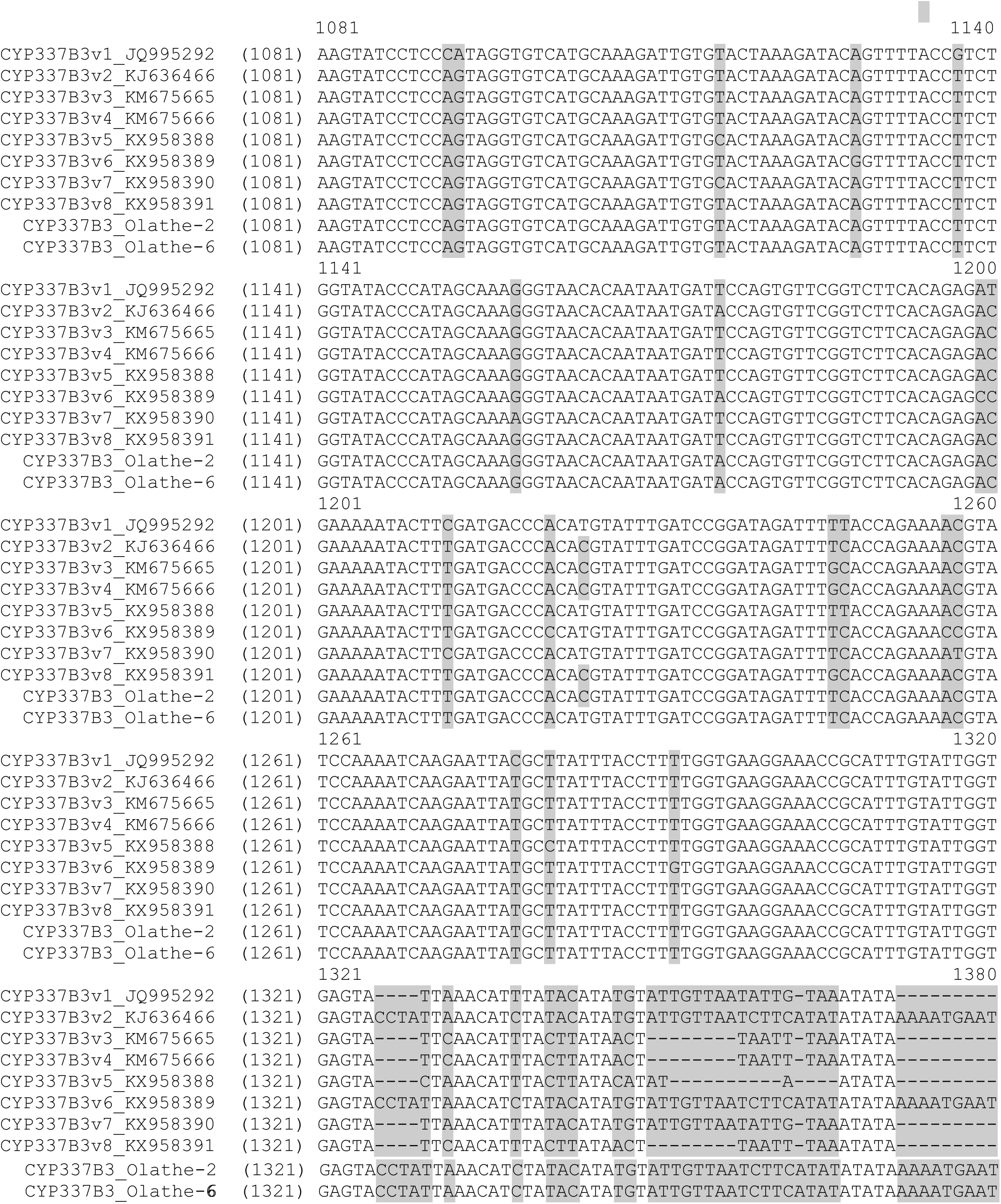
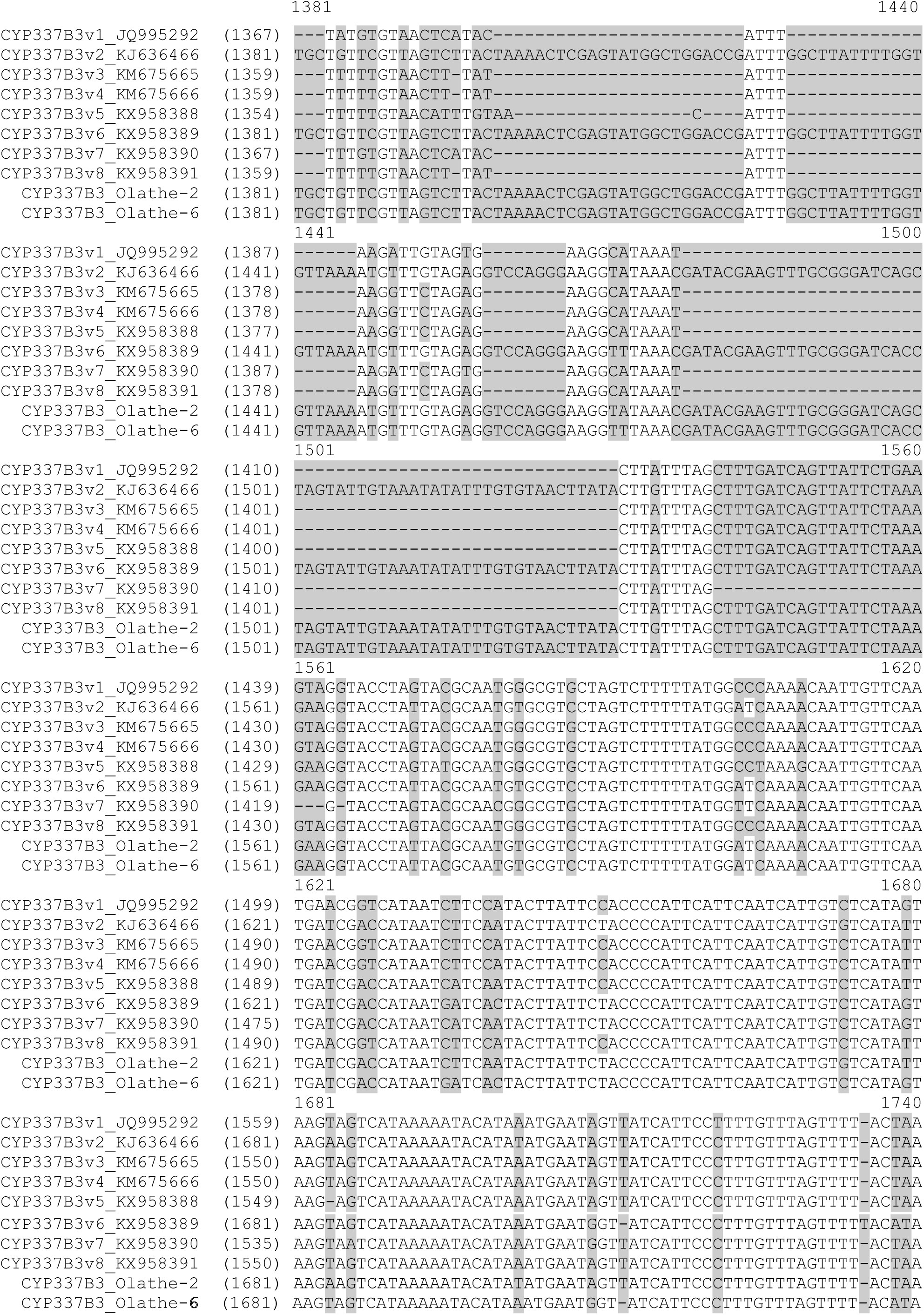
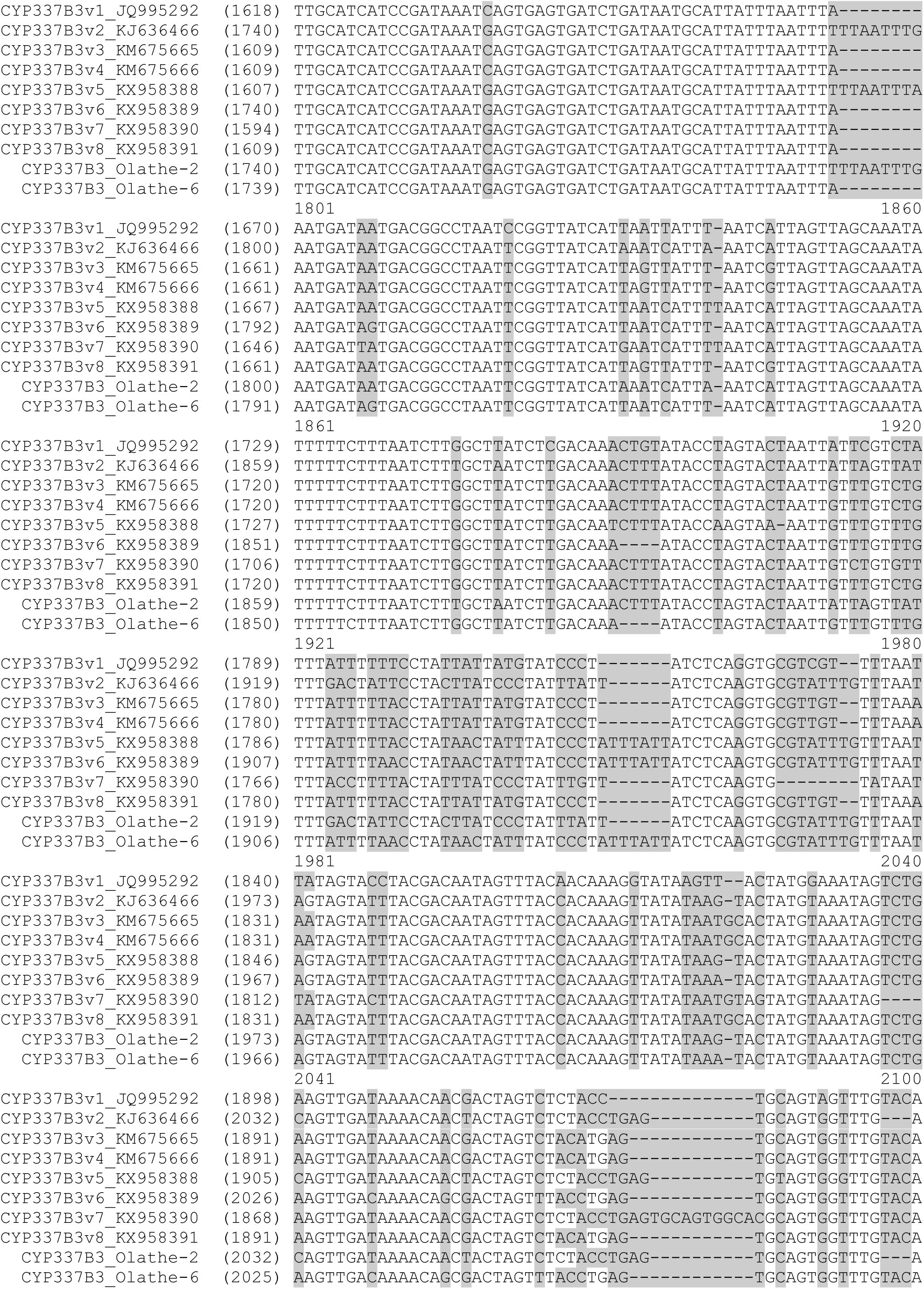
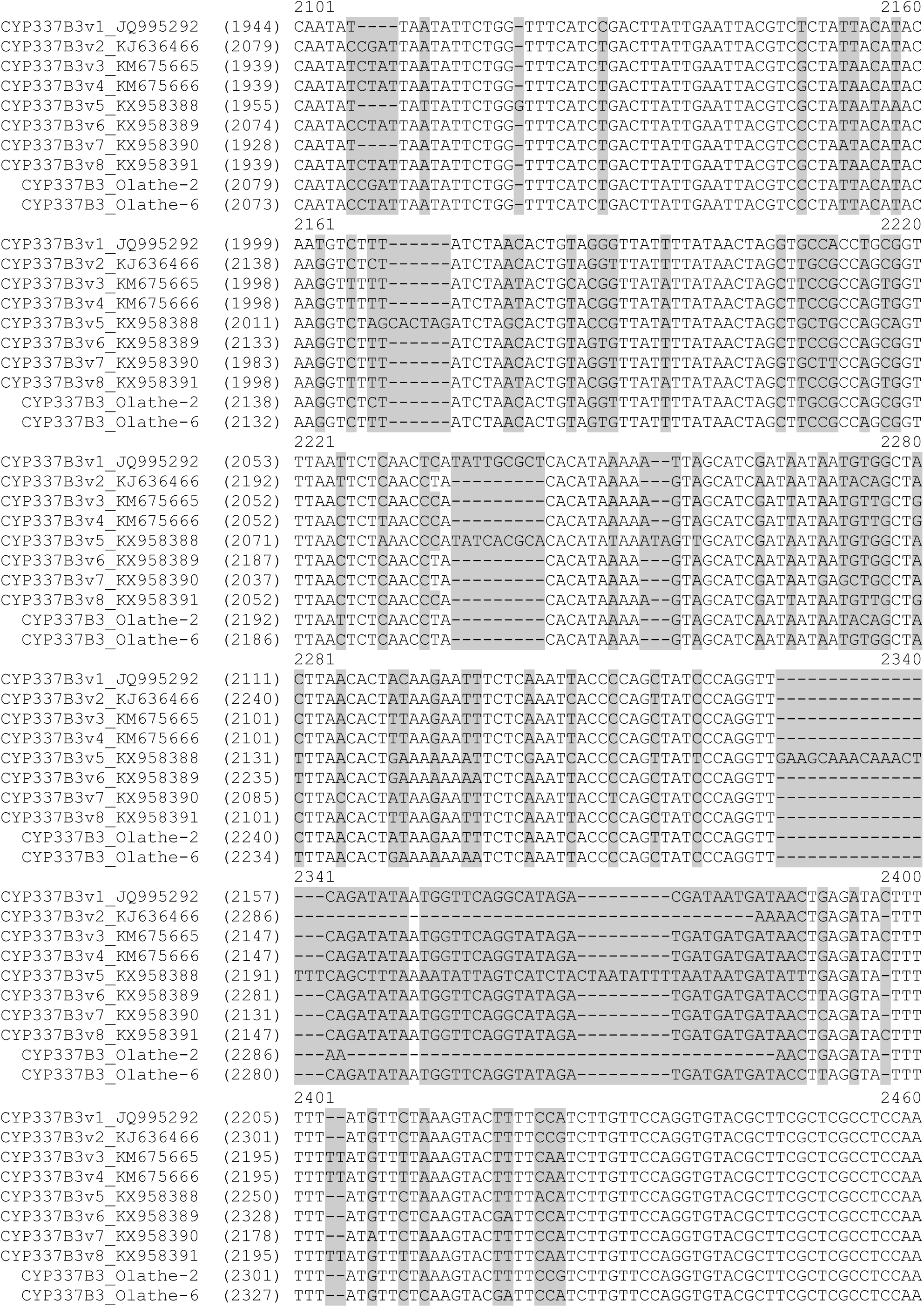
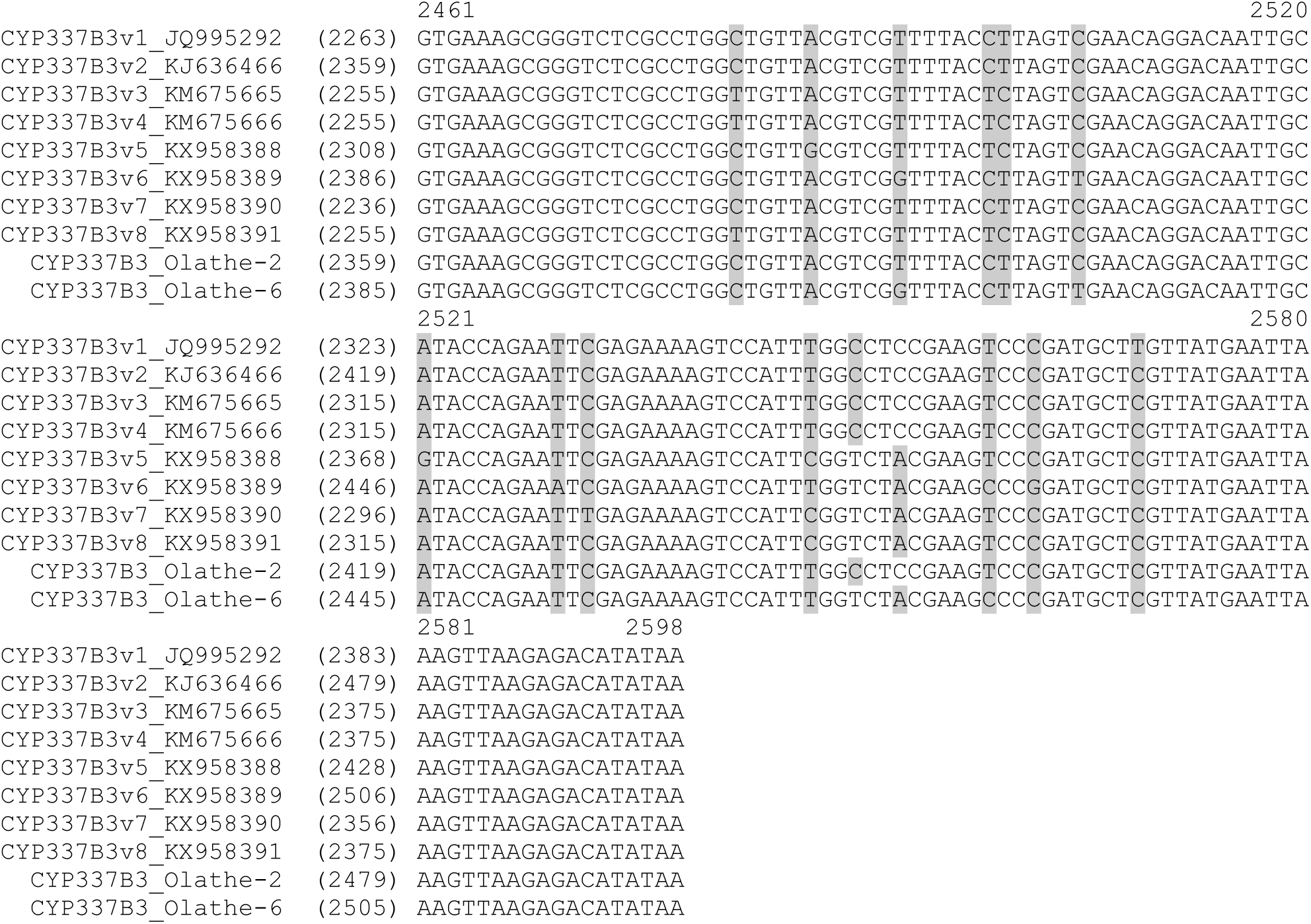
Alignment of reference genomic DNA sequences of cytochrome P450 337B (CYP337B3 gene alleles v1 through v8 and CYP337B3 alleles identified from Olathe, CO population of *Helicoverpa zea*. Identical and conserved nucleotide positions are shown in black text and nucleotide polymorphisms, insertions, and deletions are shown with grey background. Alignment gaps are indicated with a hyphen (-).

**Figure S5.**
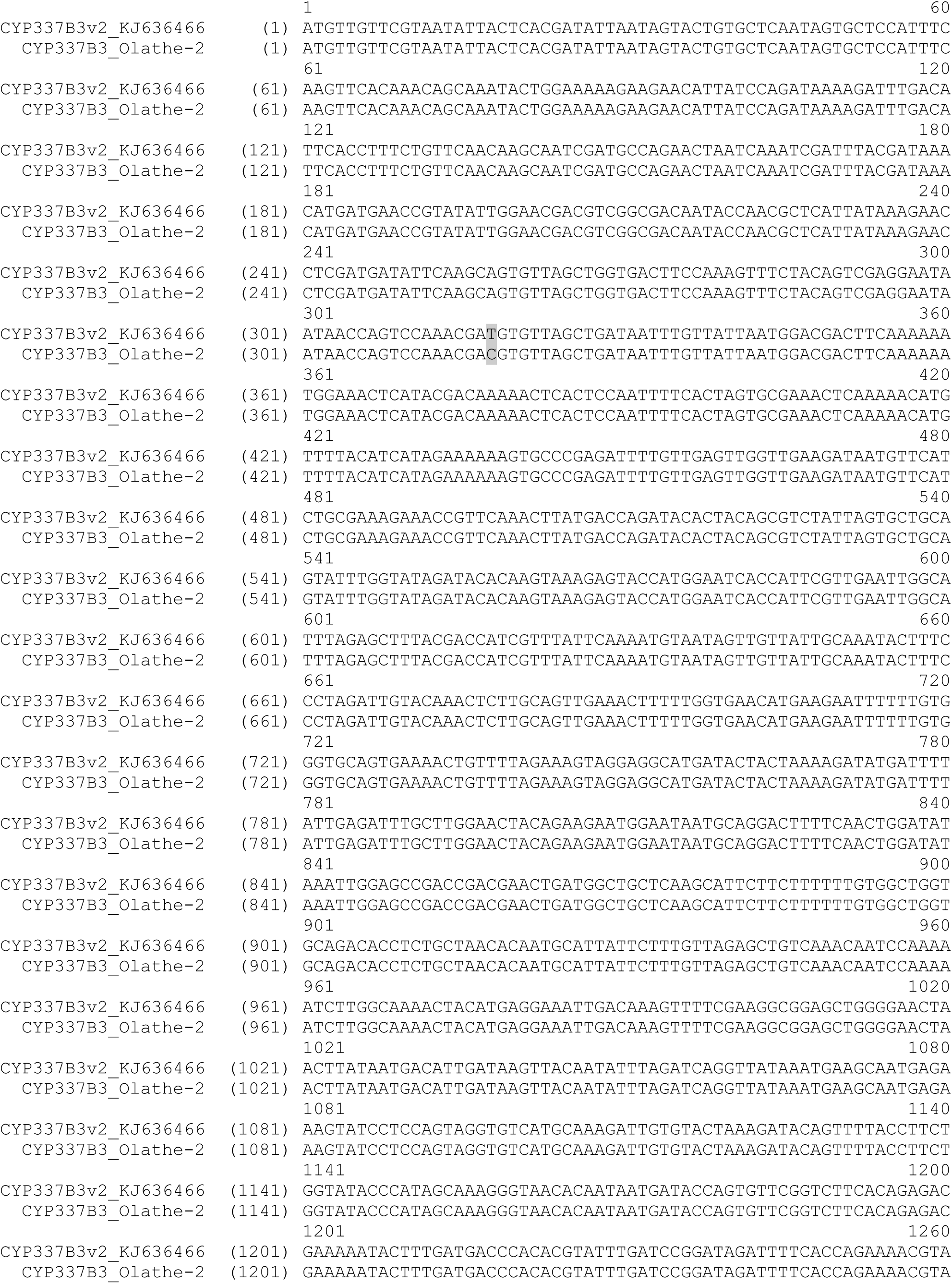
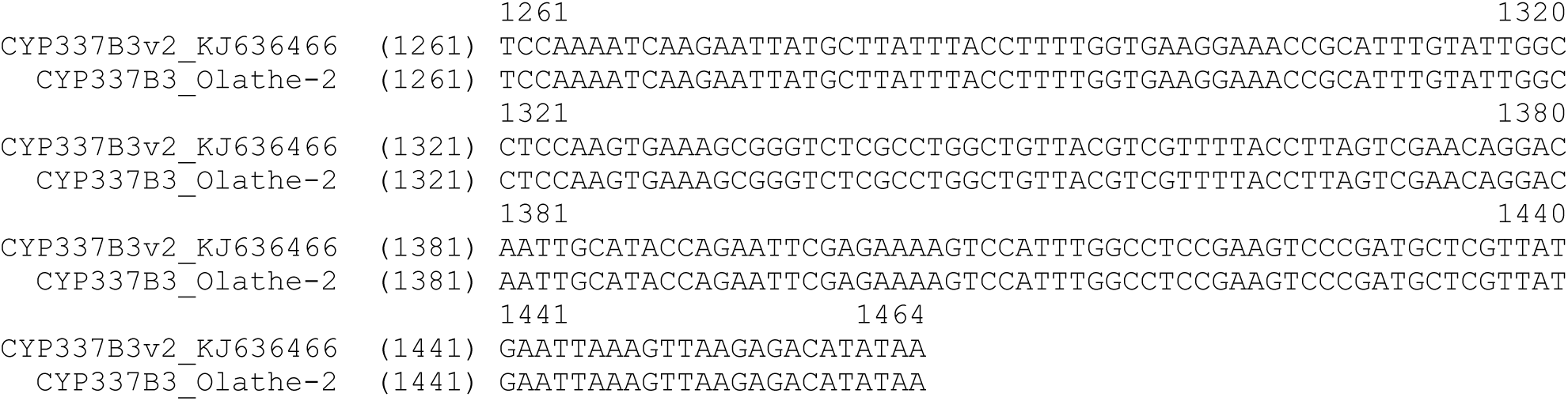
Alignment of the mRNA reference sequences of cytochrome P450 337B3 allele v2 (KJ636466.1) and CYP337B3 allele v2 identified from Olathe, CO population of *Helicoverpa zea*. Identical and conserved nucleotide positions are shown in black text and nucleotide polymorphisms, insertions, and deletions are shown with grey background. Alignment gaps are indicated with a hyphen (-).

**Figure S6.**
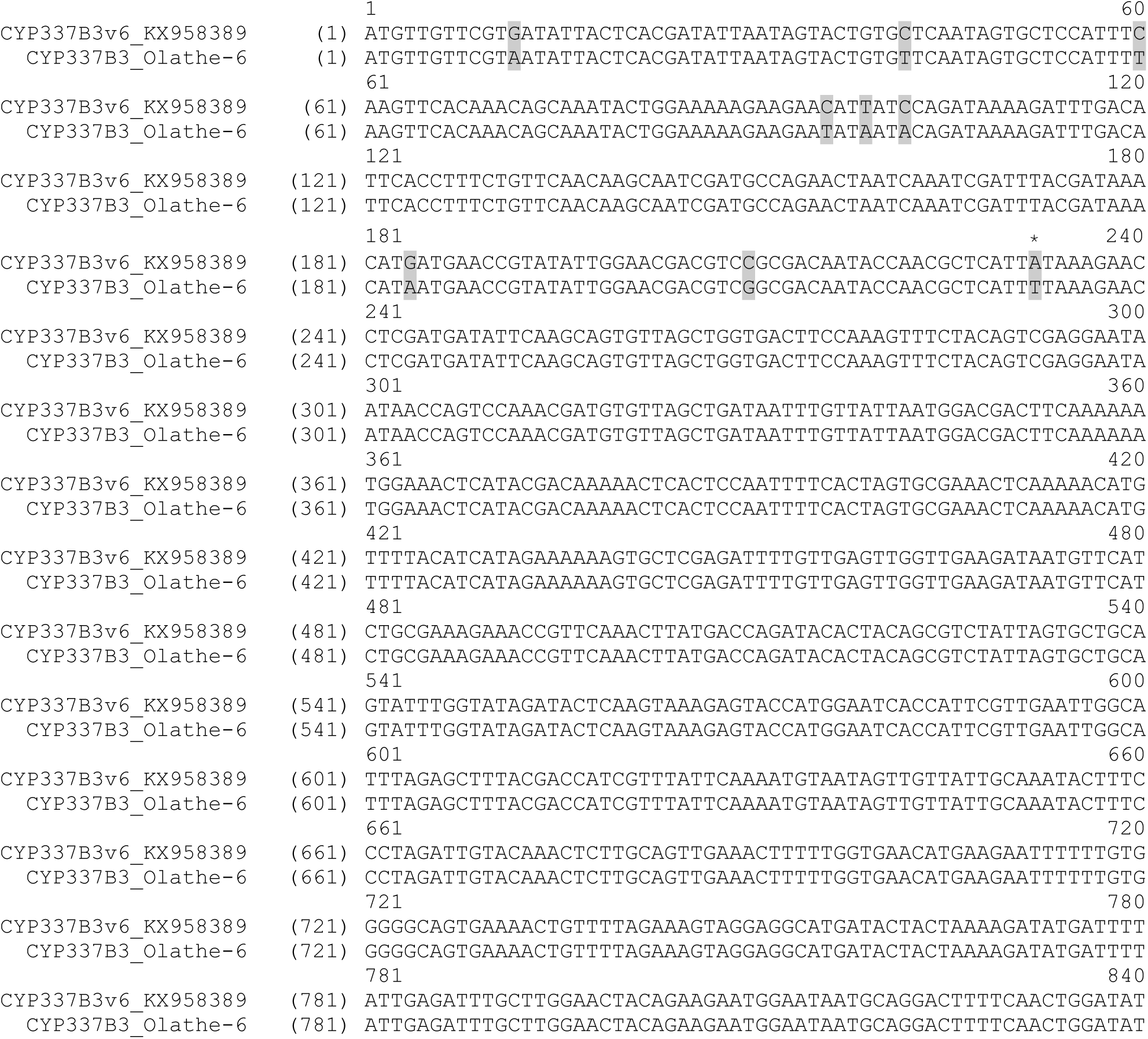
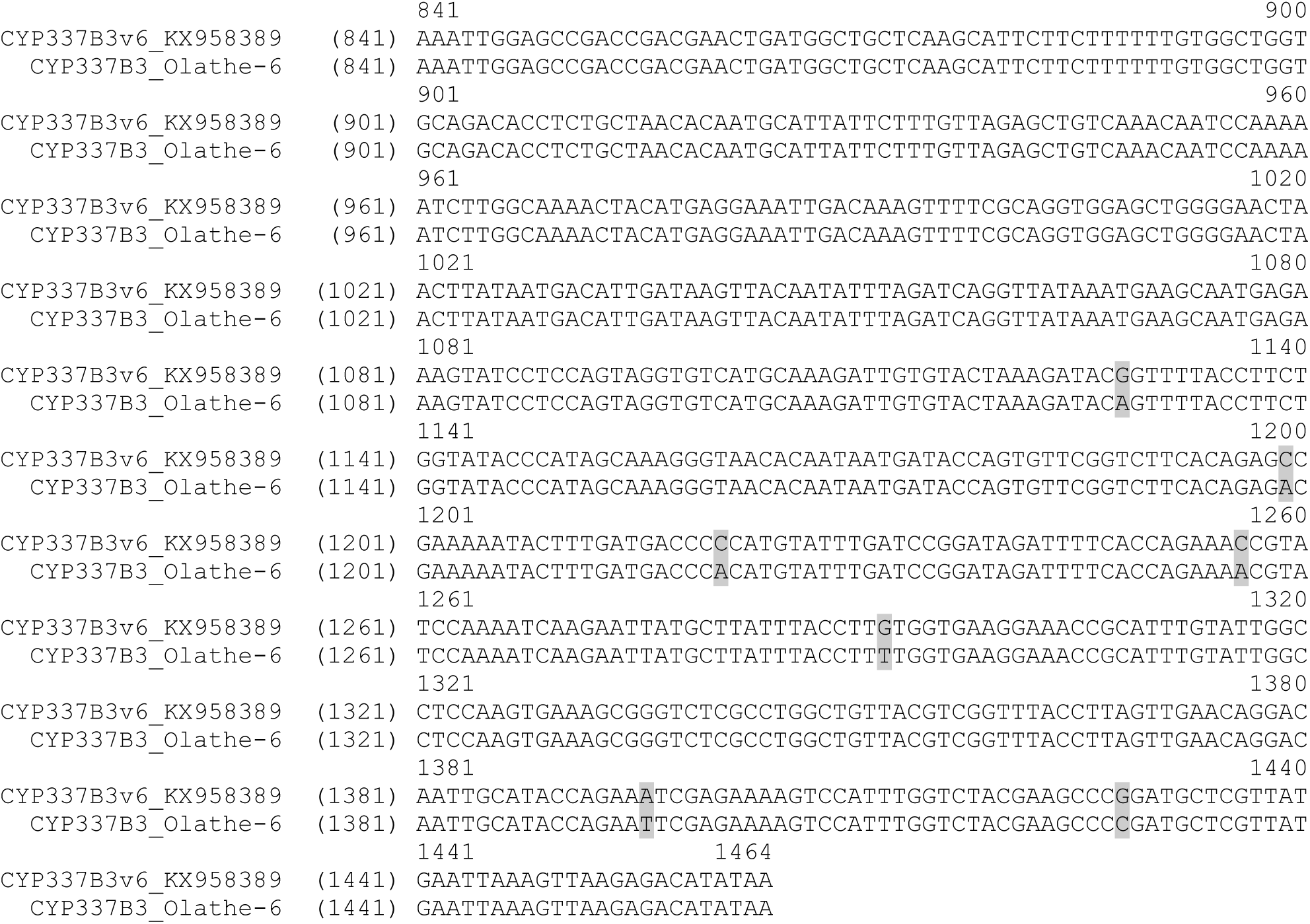
Alignment of the mRNA reference sequences of cytochrome P450 337B3 allele v6 (KX958389.1) and CYP337B3 allele v6 identified from Olathe, CO population of *Helicoverpa zea*. Identical and conserved nucleotide positions are shown in black text and nucleotide polymorphisms, insertions, and deletions are shown with grey background. Alignment gaps are indicated with a hyphen (-). A to T transversion at the nucleotide position 232 leading to the amino acid substitution I78L is marked with an asterisk (*).

